# Frequency-dependent phase entrainment of cortical cell types during tACS: converging modeling evidence

**DOI:** 10.1101/2023.12.15.571874

**Authors:** Gaugain Gabriel, Mariam Al Harrach, Maxime Yochum, Fabrice Wendling, Marom Bikson, Julien Modolo, Denys Nikolayev

## Abstract

**Background:** Transcranial alternating current stimulation (tACS) enables non-invasive modulation of brain activity, holding promise for clinical and research applications. Yet, it remains unclear how the stimulation frequency affects various neuron types.

**Objective:** To quantify the frequency-dependent behavior of key neocortical cell types.

**Methods:** We used both detailed (anatomical multicompartments) and simplified (three compartments) single-cell modeling approaches based on the Hodgkin–Huxley formalism to study neocortical excitatory and inhibitory cells under various-amplitude tACS frequencies within the 5–50 Hz range at rest and during basal 10 Hz activity.

**Results:** L5 pyramidal cells exhibited the highest polarizability at DC, ranging from 0.21 to 0.25mm and decaying exponentially with frequency. Inhibitory neurons displayed membrane resonance in the 5–15 Hz range with lower polarizability, although bipolar cells had higher polarizability. Layer 5 PC demonstrated the highest entrainment close to 10 Hz, which decayed with frequency. In contrast, inhibitory neurons entrainment increased with frequency, reaching level akin to PC. Results from simplified models could replicate the phase preferences, while amplitudes tend to follow opposite trends in PC.

**Conclusion:** tACS-induced membrane polarization is frequency-dependent, revealing observable resonance behavior. This finding motivates further experimental studies of cell-specific frequency-dependent membrane responses to weak electric stimuli. Whilst optimal phase entrainment of sustained activity is achieved in PC when tACS frequency matches the activity, inhibitory neurons tend to be entrained at higher frequencies. Consequently, this presents the potential for precise, cell-specific targeting.

- Single-cell models of frequency response to tACS for a wide variety of neurons
- Simplified and morphologically accurate models were compared
- Pyramidal cells (PC) show resonant-like response close to intrinsic firing frequency
- Entrainement of GABAergic neurons increases with the frequency of tACS
- Membrane polarization resonance in GABAergic neurons within the 5–15 Hz range

## INTRODUCTION

Neocortical neuronal network oscillations are fluctuations in the extracellular potential that can be recorded as local field potentials and/or electroencephalographic signals [1]. These oscillations are tightly linked to behavior and cognitive processes [2]. In the case of neurological diseases, the extracellular potential exhibits alterations in those neural activity oscillations [3]–[6], motivating the development of brain stimulation techniques able to normalize those alterations to relieve the associated symptoms. Among the considered technologies, transcranial alternating current stimulation (tACS) is a portable, safe, and non-invasive brain stimulation technique [7] that interacts with brain activity in a phase-entrainment manner, depending on the stimulation frequency [8]– [10]. Therefore, tACS holds great potential as a therapeutic strategy for neurological disorders.

The effect of weak (on the order of 1 V/m), induced alternating EF on neural activity has been experimentally studied in vitro and in vivo in small animal models [11]–[15] as well as in primates [16], [17]. These studies demonstrated that a low alternating EF entrains neural activity, generally at the stimulation frequency. Neural entrainment of a set of interconnected neurons could synchronize activity, increasing the target network activity at stimulation frequency [18]. Although entrainment seems to be frequency-dependent, the relationship between stimulation frequency and entrainment efficiency is not straightforward. While some studies demonstrated higher neural entrainment at low frequencies [11], [12], others indicated that stimulation at the endogenous frequency was the most efficient, supporting an Arnold tongue behavior [14], [15]. This behavior was further corroborated by numerical modeling [12], [19], [20]. However, this dependency was not consistently observed experimentally and another study reported that tACS could compete with endogenous activity, being less effective when applied at the intrinsic frequency [21]. Other results provided evidence of a non-linear dose-response to 140 Hz tACS with a switch from an inhibitory to excitatory effect at 0.4 V/m, suggesting a higher sensitivity of inhibitory neurons at low EF intensities and excitatory cells at higher EF intensities [22]. Conversely, tACS effects were attributed almost exclusively to excitatory pyramidal cells. Consequently, the underlying cellular-level mechanisms of tACS are still debated, and its frequency-dependent effects are only partially understood.

In this context, computational modeling is a promising solution to uncover tACS mechanisms and explain certain experimental results, enabling testing of wider parameter sets that are inaccessible in *in vivo* conditions. The great majority of computational modeling studies used simplified cells either to study single-cell or network tACS effects [19], [23]. While this simplified approach is useful to save computation time (enabling network simulation that would be unsuitable with more realistic models), it requires calibrations to properly account for EF coupling since morphology is not accounted for. A recent study used realistic neuron models to investigate the 10Hz–tACS effect at the single-cell level with multiple reconstructed neuron morphologies including full axonal tree and myelin inclusion [24]. This study found that pyramidal cells, especially those from layer 5 (L5 PC) were the most sensitive to tACS, but also that inhibitory neurons could be more sensitive to tACS entrainment than L6 or L2/3 pyramidal cells. Nevertheless, the cell models used were adapted independently from activity measurements, which could lead to ”unnatural” behavior [25].

In this study, we used realistic modeling of single cells, with a broader variety of inhibitory cells, to assess their sensitivity to tACS parameters (frequency, intensity). Two levels of physiologically relevant models of neurons and interneurons were used: “detailed” morphologically reconstructed models, and “simplified” 3-compartments models. First, we studied the membrane polarization spectral behavior of each cell’s membrane for the detailed models. Then, we applied a realistic synaptic inputs set to generate a middle alpha band activity of 10 Hz to investigate the neural entrainment to tACS waveform using both models.

## MATERIALS AND METHOD

### Detailed neuron models

Morphologically reconstructed neuron models proposed in [26] were used as realistic models of neuronal geometry. These models were originally reconstructed from images of juvenile rat somatosensory cortex neurons. The associated reconstructed neurons were then segmented into compartments that are compatible with NEURON [27]. This enables the standard simulation of neuronal activity by solving the cable equation for each compartment:

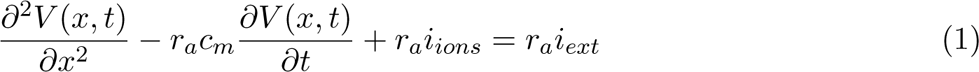

where V (x, t) is the membrane potential, r*_a_* the axial membrane resistance, c*_m_*its specific capacitance, i*_ions_* the ionic currents due to the different membrane ion channels, and i*_ext_* is an exterior current (i.e., due to external stimulation). A set of 13 Hodgkin-Huxley type channels were used to fit data from electrophysiological recordings after removing the reconstructed axon to replace it with a two-compartments straight axon (see [26]). Axonal trees were available for each cell.

However, neuron channel parameters were fitted with a straight axon, and since it is known that neuron activity is dependent on its geometry [28], straight axons were maintained to preserve the original measured activity. Our strategy differs from Tran *et al.* (2022) [24], who used a myelinated full axonal tree built in [25]. [25]. Here, in-house python scripts were developed to control all simulations.

Among the various morpho-electrical cell types available, a specific subset was chosen for the sake of computational cost. This subset included layer 5 thick tufted with early bifurcation pyramidal cells (L5 TTPC2, for comparison with [24]), layer 2/3 pyramidal cells (L2/3 PC) and a set of layer 4 inhibitory cells which best represents vaso-intestinal peptide (VIP), somatostatin (SST) and parvalbumin (PV) expressing interneurons, as depicted Figure 1.B (please refer to Table S.2.1 for a comprehensive list).

**Figure 1:**
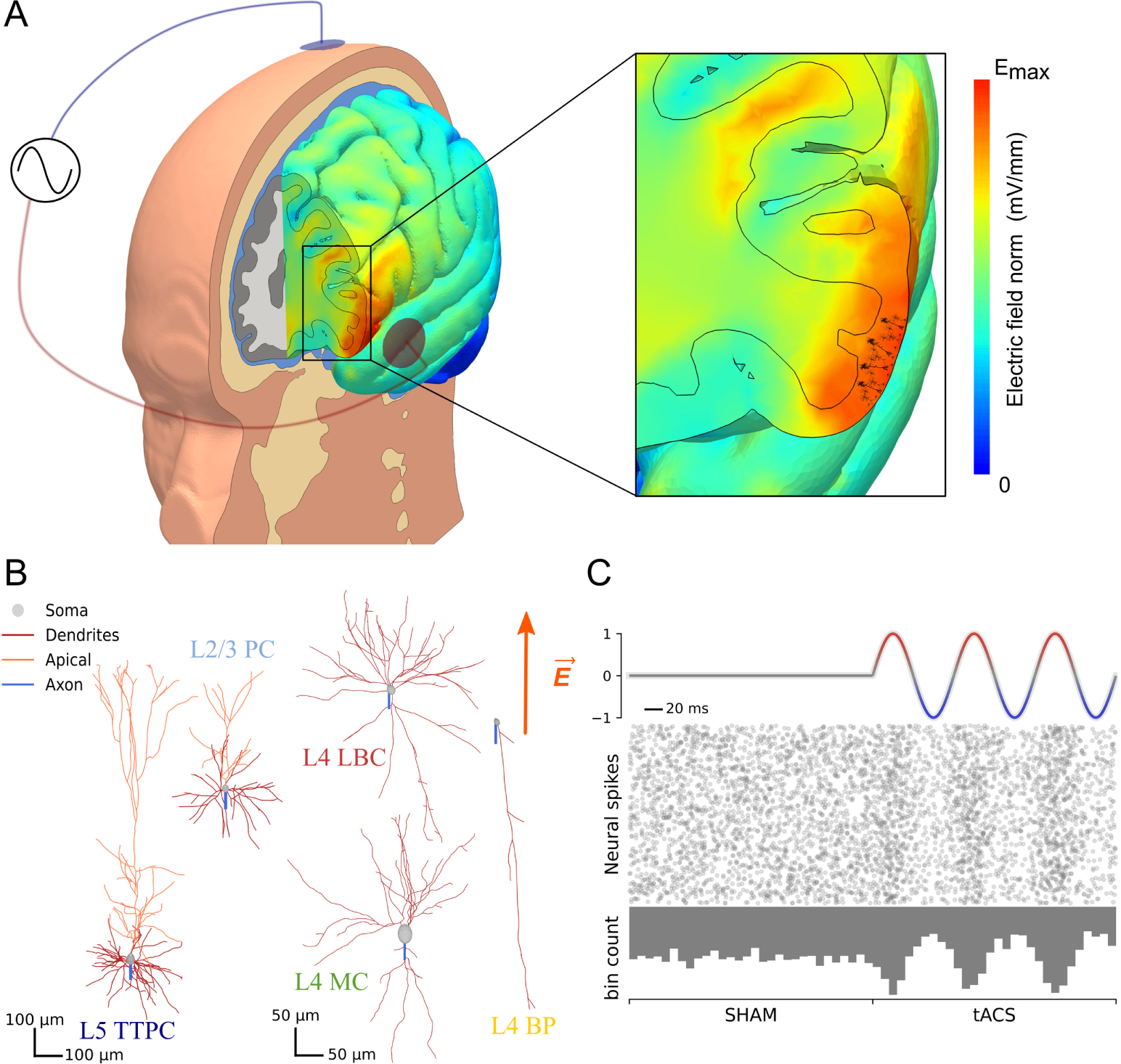
A: Schematics of a bipolar tACS montage with two 2-cm diameter electrodes. The five tissues commonly considered for tACS dosimetry are depicted along with the resulting electric field in the brain. A zoomed-in view of the example targeted area where the maximal electric field and sample studied neurons are depicted. B: Subset of considered neuron morphologies with L5 TTPC and L2/3 PC cells, and three L4 inhibitory neurons as LBC (PV), BP (VIP), and MC (SST) cells. EF orientation is depicted as E^⃗^. C: Neuronal activity affected by the EF through phase entrainment. The raster plot shows spiking events during the stimulation depicted in the first row. In the absence of stimulation (SHAM), spikes are randomly distributed (left side). During tACS, spiking events are preferentially distributed in the positive phase of tACS (darker raster plot), and less during its negative phase (lighter raster plot). The associated bin counts depicted in the lower portion of C highlight further this trend, with the bin count following the tACS waveform.

### Simplified single cell models

A spatially reduced version of the multi-compartment morphological model was used in this study to represent both PCs and interneurons. This simplified model was adapted from the work of Demont-Guignard *et al.* 2009 [29]. These models encompass a greater variety of currents compared to traditional reduced models with a balance between physiological accuracy and computational efficiency. In this study, we introduced a novel three-compartment model for neocortical pyramidal neurons, comprising the soma, dendrites, and axon initial segment (AIS), as illustrated in Figure 2. The Hodgkin–Huxley formalism was used to simulate the membrane potential of each compartment [30], [31]. Using the conservation of electric charge equation presented below:

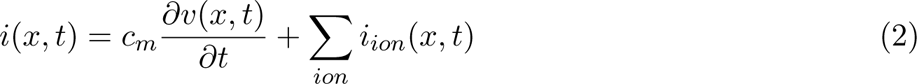

**Figure 2:**
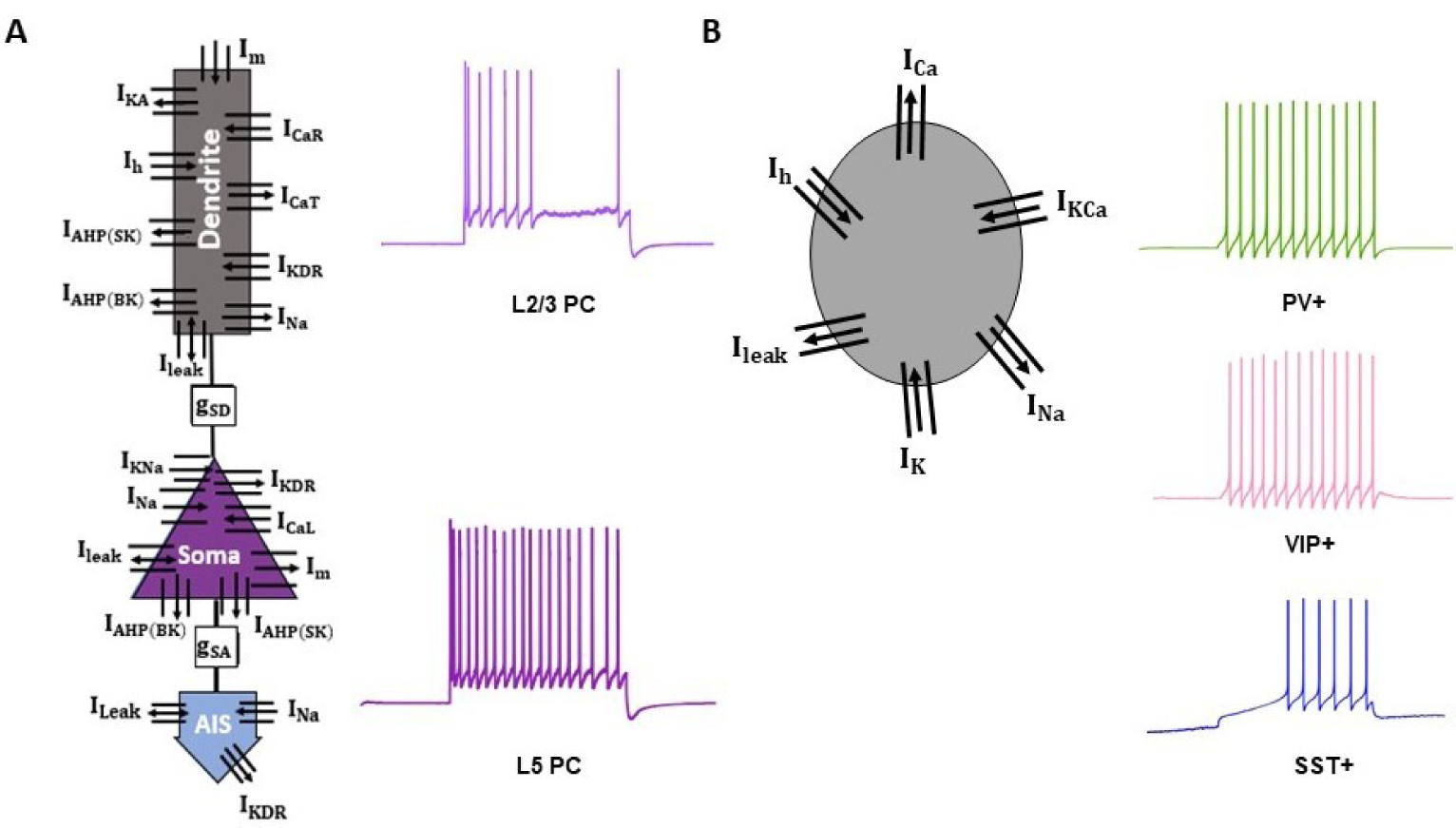
Simplified neocortical cells computational models. A. Reduced three-compartments (dendrite, soma and Axon Initial Segment (AIS)) model for pyramidal neurons (L2/3 and L5 PC). B. Single-compartment model for ParValbumin (PV+), SomatoStatin (SST+) and Vasoactive Intestinal Polypeptide (VIP+) expressing interneurons. Ionic currents were as follows: Voltage-dependent sodium current (i*_Na_*), potassium current (i*_K_*), potassium delayed-rectifier current (i*_KDR_*), calcium-dependent potassium currents (i*_AHP_* ), muscarinic current (i*_m_*), L/R/T-type calcium currents (i*_CaL_*, i*_CaR_* and i*_CaT_* ), fast inactivating potassium current (i*_KA_*), hyperpolarization-activated cationic current (i*_h_*), sodium activated potassium current (i*_KNa_*), and calcium-activated potassium current (i*_KCa_*).

where i(x, t), v(x, t) and i*_ion_*(x, t) are the injected input current, membrane potential, and ionic currents of compartment x, respectively. The coupling of the soma/dendrites compartments and Soma/AIS was made through the conductances g*_SD_* and g*_SA_*, respectively (Figure 2.A). The ionic currents were specific to each compartment and cell type (Figure 2), and were described by:

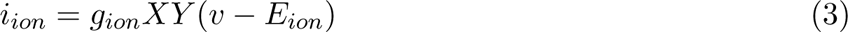

With g*_ion_* the ionic conductance, E*_ion_* the reversal potential, and XY the probability of channel opening. Ionic channel parameters’ values were adjusted to replicate the firing rate obtained through electrophysiological recordings.

### Synaptic input

The simplified model included both glutamatergic α-amino-3-hydroxy-5-methyl-4-isoxazolepropionic acid (AMPA) and N-methyl-D-aspartate (NMDA) and GABAergic synapses as in [29]. Only excitatory synapses were used to generate physiological activity. For realistic models, the original statistically reconstructed set of synapses for each individual cell provided by the Blue Brain Project (available in NMC-portal) was used for reproducibility. Synapse parameters were fitted for both excitatory (AMPA/NMDA) and inhibitory (GABA A/B) synapses using a Tsodyks-Markram-based model [32]. For both models, presynaptic spike trains were generated using a stochastic Poisson distribution, to produce a mean firing rate at 10 Hz. To achieve reproducible spiking activity around 10 Hz in the realistic model, which has multiple synapses, excitatory and inhibitory synaptic weights were scaled inversely (see Table S.2.1 for the values and corresponding firing rates).

### tACS modeling

For the realistic model, extracellular stimulation due to tACS was achieved using the extracellular mechanism included in the NEURON simulator [27] together with the xtra mechanism to control the stimulation waveform. The quasi-uniform approximation [33] was used, and the electric field was set to be uniform and oriented in the y-direction, which was the somato-dendritic axis of the pyramidal cells. The resulting extracellular potential was calculated as e*^extra^*(t) = E_0_ *·* a(t) *·* x*_i_*, where e*^extra^*(t) is the extracellular potential at the i*^t^*h compartment at time t, x*_i_* its coordinate, E_0_ the unit electric field vector which was [0, 1, 0] here, and a(t) the stimulation waveform at time t.

To investigate membrane polarization due to tACS, simulations were performed with initialization to the steady state and no external inputs. We investigated the [0, 50] Hz tACS frequency range with an increment of 0.25 Hz, while simulation time was set to 4 s if the frequency was below 5 Hz, and 2 s for higher frequencies since this duration was sufficient to reach steady-state.

We investigated tACS effects in both models by defining the waveform function as a(t) = A *·* sin(2πft), where A is the EF amplitude and f is the tACS frequency. We explored a range of intensities from 0 (no stimulation) to 10 V/m to identify the threshold for significant entrainment to the EF for each single cell, while acknowledging that this upper limit of EF value is higher than can be achieved during standard tACS protocols. However, our goal was to identify the cells that are individually the most sensitive to tACS (lower threshold), and to identify a threshold for entrainment effects, justifying our approach.

To model the impact of tACS on the activity of single cells in the simplified model, the “lambda-E” was used [34], [35]. This model states that for weak and uniform electric fields, the membrane polarization of all compartments is linear with the stimulation current amplitude. Accordingly, the membrane polarization induced by the EF at a compartment is described as the dot product of EF vector and a length vector representing the coupling constant [34]:

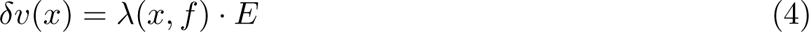

Values of the length vector for the different cell types simulated in this study are obtained from [34]. for DC (see Table 1). λ were scaled over the frequency spectrum using normalized values obtained with the realistic cells, since membrane polarization is frequency dependent [36].

**Table 1:** Lambda (λ) values for the five cell types simulated using the simplified computational model of single cells. These values represent the norm of the λ vector in mV, and are adapted from [34].

|  | PC |  |  | VIP | SST | PV |
| --- | --- | --- | --- | --- | --- | --- |
|  | Soma | Dendrites | AIS |  |  |  |
| L2/ 3 | 0.032 | -0,120 | 0.032 | 0.200 | -0.080 | 0.090 |
| L4 | - | - | - | -0.025 | -0.070 | 0 |
| L5 | 0.137 | -0.071 | 0.137 | -0.085 | -0.080 | 0 |
| L6 | 0.117 | -0.046 | -0.117 | -0.100 | -0.060 | -0.070 |

The simulation of tACS effects at the single cell level was organized as follows: 2 minutes of pre-stimulation activity or control activity, followed by 4 minutes of tACS (Figure 1.C). A control simulation of 6 minutes with no tACS was performed for comparison. The Poisson noise distribution was seeded in the simulation to ensure the same external input was sent to the neuron for every stimulation parameter pair.

### Phase entrainment quantification

The spike events from the simulated 6 minutes of activity (4 minutes of tACS) were extracted with the voltage trace from the first axonal segment (AIS). Subsequently, the firing rate (FR) was calculated during and before tACS to quantify the effect of tACS on FR, which was not expected to be significantly affected [24], [37]. To quantify neural entrainment due to tACS, we computed the phase locking value (PLV) [38] defined as:

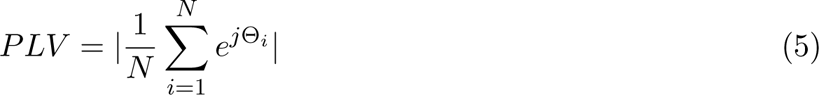

where N is the number of spike events, and Θ*_i_* the phase of the tACS waveform at which the i*^th^* spike occurs. This metric quantifies the spike synchrony relative to tACS with a value ranging from 0 (no synchronization) to 1 (perfect synchronization). To assess deviation from a circular uniform distribution, the Rayleigh test was used [39]. Polar plots were used for visual representation of these quantitative values, since a preferred phase direction is associated with more bin counts in an angular direction.

## RESULTS

### Cell frequency dependent sensitivities to electric field

By examining the frequency dependence of all considered cell membrane polarization due to tACS using the detailed model, we found different spectrum profiles for PC, SST, PV, and VIP. The re-sulting spectra are depicted in Figure 3.A. L5 PCs had the highest polarization length for a direct current (DC) stimulation (0.224-0.255 mm), with a decreasing profile when frequency increased. The profile was similar for L6 PCs, with the lowest values for DC (0.15-0.18 mm), but in a similar range from 20 Hz to 50 Hz (0.05-0.10 mm). L2/3 PCs polarization had fewer variations across the considered spectrum, with the lowest polarization from all PCs. The second highest polarization length cell type was found to be BP, in particular with bNAC dynamics, while cAC dynamics decreased PLV. Frequency dependence was shared for both dynamics, and strongly differed from PCs since it monotonically increased from DC to 10 to 15 Hz to a maximum value, then monotonically decreased. SST and PV cells had a similar range of polarization length (0.005-0.07 mm), with the lowest value at DC for most cells. In the MC, the polarization length spectrum monotonically increased up to 5 Hz, and slowly decreased in the remaining frequency range. However, values were strongly variable across similar cell types. In the case of PV cells, since more than one cell type and dynamics were tested, the trend varied across cell types and dynamics.

**Figure 3:**
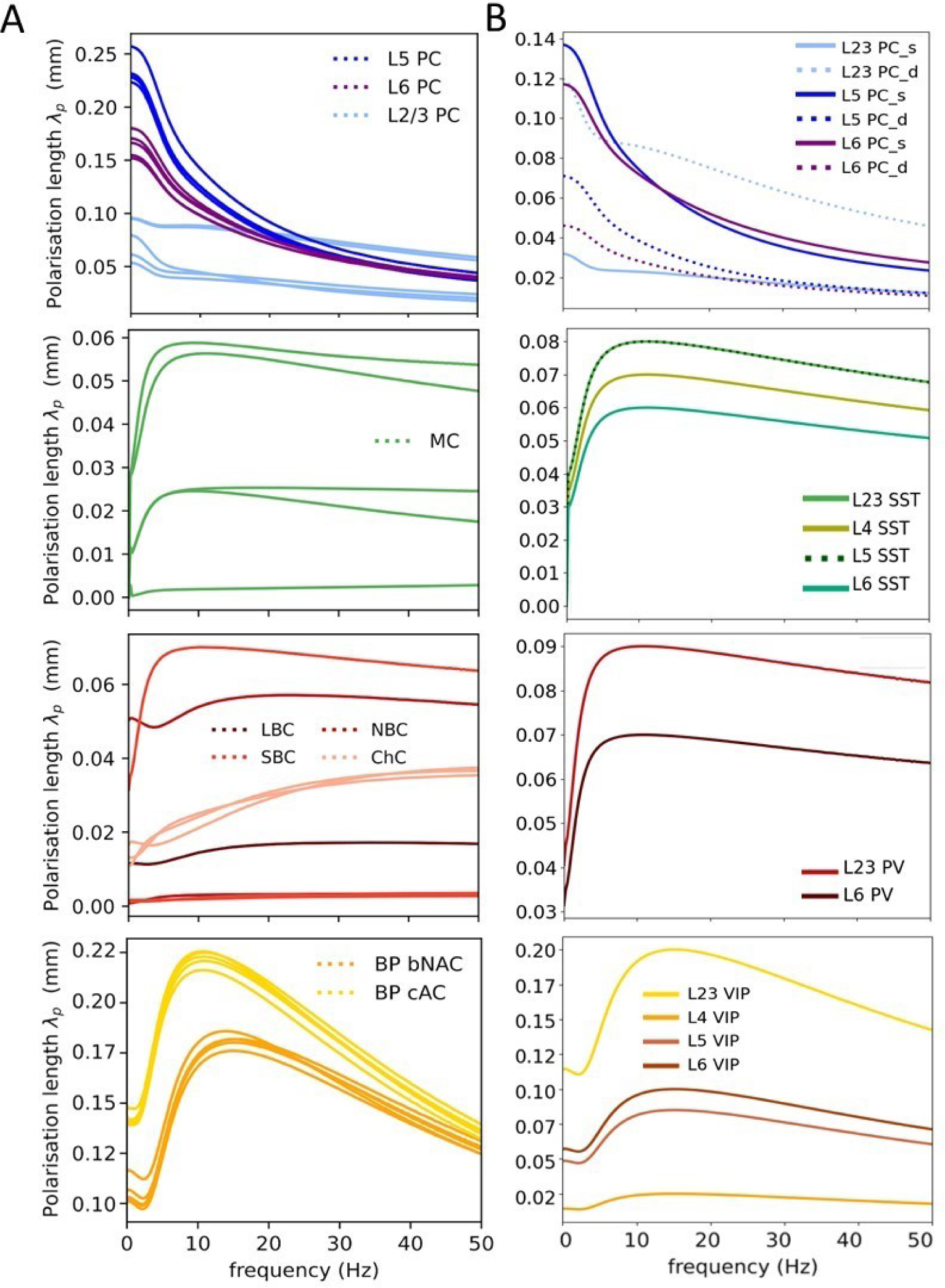
tACS frequency dependence of polarization length for all tested cells using the detailed (A) and simplified models (B). Cell types from top to bottom are as follows: PCs, SST, PV, and VIP. For the simplified model, λ values of soma and dendrite compartments are depicted for the PCs. The AIS compartment was considered to have the same λ values as for the soma.

In the case of the simplified model, the modeling of tACS effects followed the lambda-E model, with lambda values obtained from the steady-state change in membrane potential induced by a 1 V/m electric field [34]. However, it is important to note that these values were originally obtained in the context of transcranial Direct Current Stimulation (tDCS), and do not account for the frequency-dependent nature of tACS. To address this and to adapt the simplified model from the detailed model, and considering the distinct morphological types of cells, we used the polarization length profiles computed using the detailed model (refer to Figure 3.A) to adapt lambda values for the lambda-E model. In this configuration, we assumed that the variation profile was the same for all compartments. Figure 3.B illustrates polarization length variations with respect to tACS frequency in the soma and dendritic compartments for L2/3, L5, and L6 PCs as well as for VIP, SST, and PV cells, respectively, for all neocortical layers. We can observe that, among PCs, the dendrites of L2/3 PC have the highest polarization length for tACS from 10 to 50 Hz. However, in the case of DC stimulation, the soma of L5 and L6 PC had the highest polarization length. For interneurons, SSTs depicted similar variations throughout the layers. L23 PV and VIPs had the highest polarization as compared to other layers (Figure 3.B). If we compare the polarization profiles between the two models, we can deduce that we have a similar variation range for all considered cells except L5 PC which is twice higher in the case of the detailed model.

### Cell type specific dose-response to 5, 10, 20, and 40 Hz–tACS on phase entrainment

We examined the effect of four different tACS frequencies for each cell (5, 10, 20, and 40 Hz tACS) with increasing intensities. All investigated neurons had a generated activity around 10 Hz, as described above. No change in the firing rate (FR) was observed during tACS compared with the control simulation (see Figure S.2-4.), as expected due to the fact that the waveform is biphasic and in accordance with the literature. For all frequencies, a linear increase in entrainment to tACS frequency with EF intensity was found, as illustrated in Figure 4 for both models. This is in line with previous in vivo [36] and computational studies [24]. According to the detailed model, PC from L5 had the highest slope, followed by L6 TPC for all frequencies (Figure 4.A). However, the mean slope for the L5 PC exhibited a maximum for the 10 Hz tACS and decreased with frequency (see Figure 4.A and Table S.2.1). L6 TPC did not have a significantly different mean slope between 10, 20, and 40 Hz with a lower slope at 5 Hz. In fact, all cells but L2/3 PCs had their lowest slope at 5 Hz. An increase in slope with stimulation frequency was found for SST and PV cells, with a slope even higher than L2/3 PC at 20 and 40 Hz. Two PV and one SST cells had similar PLV slopes and ranges during 40 Hz–tACS than L5 PC during 10 Hz–tACS, as depicted in Figure 5.C & F (see Figures S2.5-8 for individual PLV plots for further comparisons).

**Figure 4:**
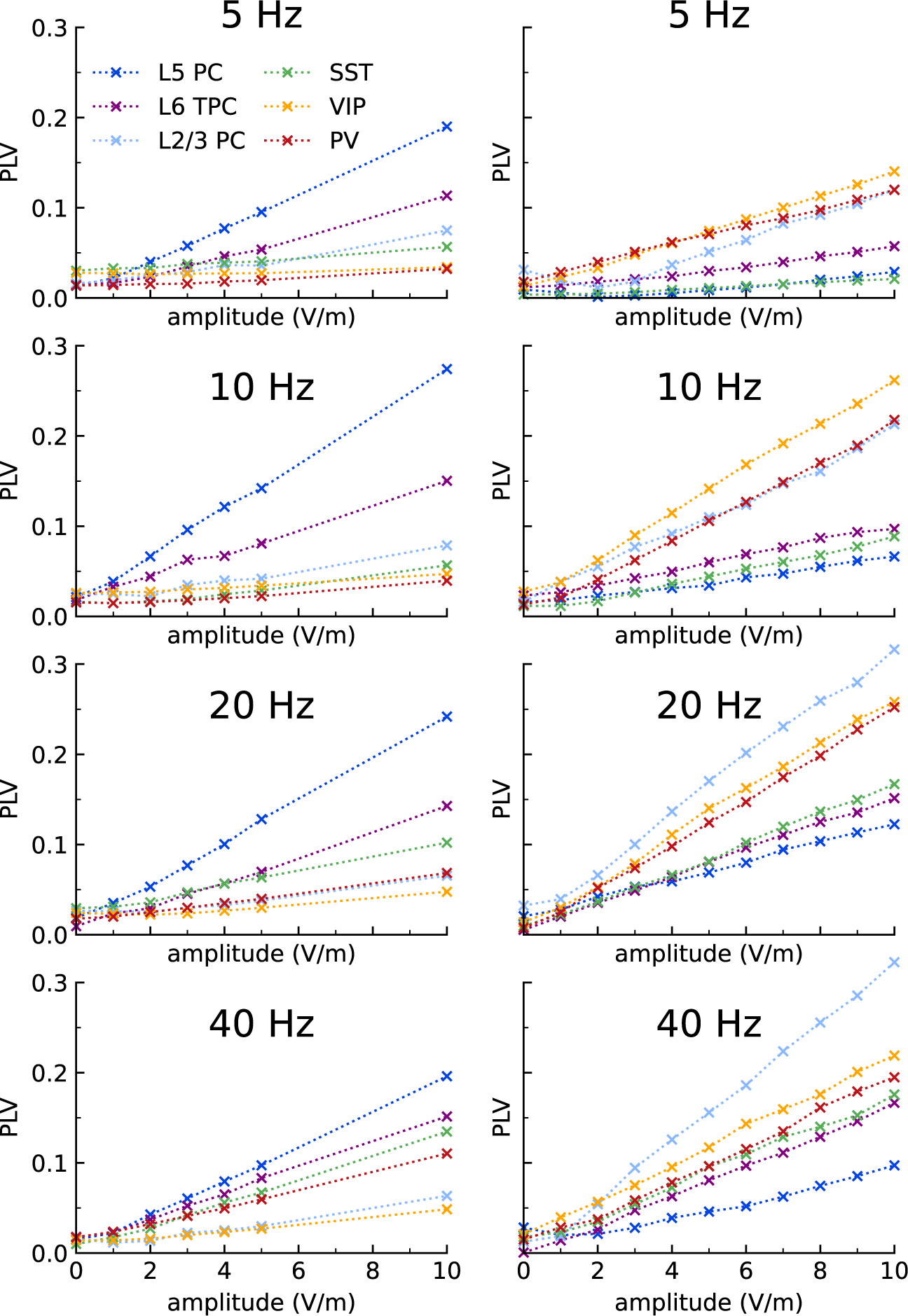
Measure of neural entrainment (PLV) at 5, 10, 20, and 40 Hz-tACS for considered cell types (L5 PC, L6 TPC, L2/3 PC, VIP, SST, and PV) using the detailed model (A) and simplified cell model (B). (A) PLV is plotted for visibility while complete plots for all cell types are available in supplementary materials (Figures S2.5-8). (B) For PCs, individual variations are plotted as for SST, PV, and VIP interneurons, the plots represent the mean variation of PLV for L23, L4, L5, and L6 cells. Detailed plots for each layer are given in supplementary Figure S1.1. were significantly entrained by 10 Hz–tACS as well as one SST cell (one MC cell) and one VIP cell (BP bNAC cell). During 40 Hz–tACS, only one PV cell was significantly entrained at 1 V/m while one L5 PC, one L6 PC, two PV, and one SST were entrained at 2 V/m. This supports preferential inhibitory targeting with 40 Hz–tACS, while excitatory cells seem more entrained at their spiking frequency. Figure 5.C shows the mean PLV polar plot and corresponding Preferred Phase (PPh) for L5 PC and L4 PV cells obtained for both simplified and detailed models. Therefore, both models result in similar PLV variations. However, one discrepancy was an offset of about 45° in the PPh phase between the two models.

**Figure 5:**
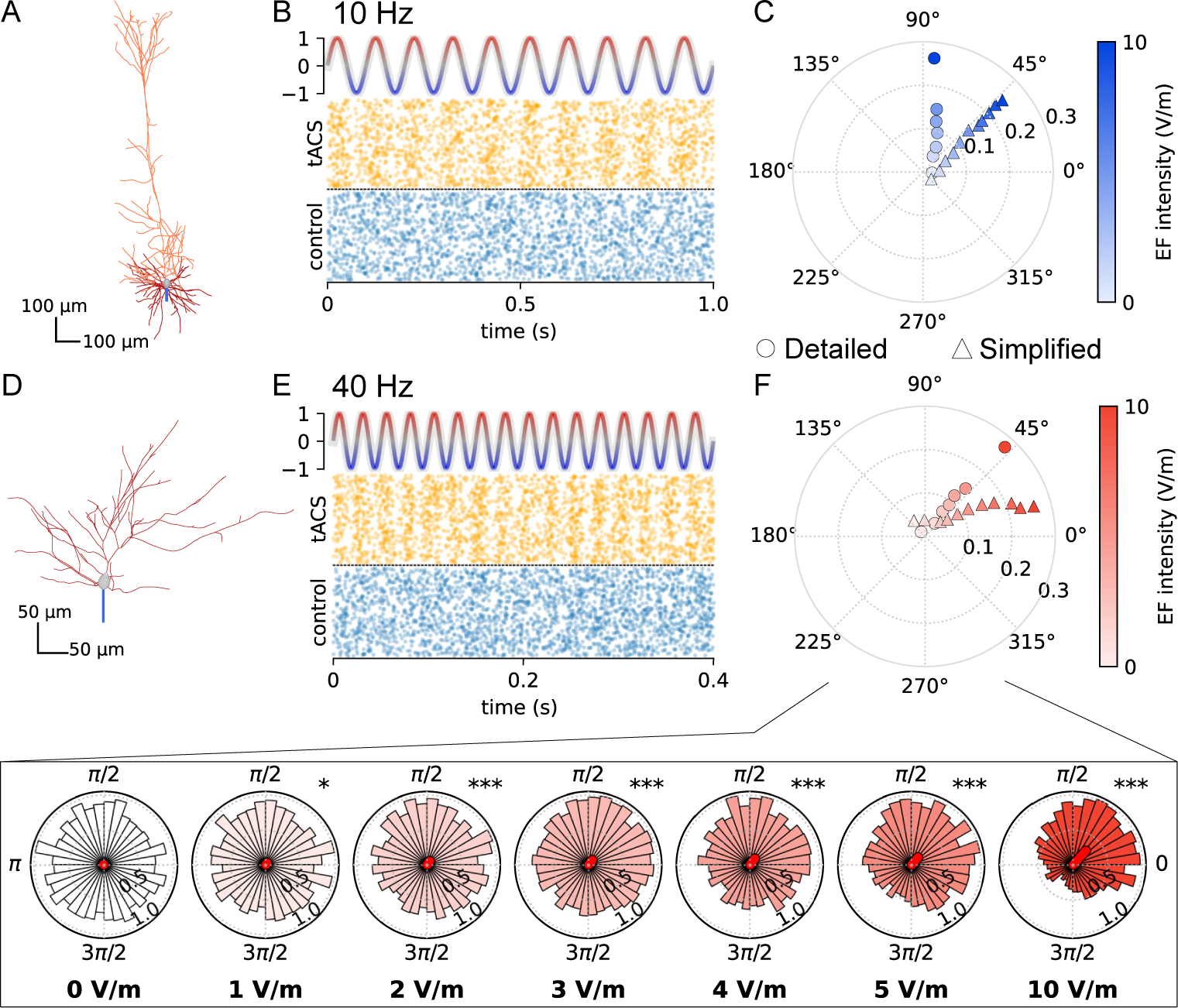
Example of a responsive cell for 10 Hz and 40 Hz tACS. A. L5 PC morphology associated with the raster plots for control and 10 V/m 10 Hz–tACS (B). C. Polar plot of the mean vector (PLV with associated phase preference) computed for using both detailed and simplified models. D-F. The same representations for a responsive L4 PV cell (small basket cell) during 40 Hz–tACS.

In the case of the simplified model, among PCs, L2/3 PC exhibited the highest entrainment levels (Figure 4.B). All PCs had an increasing PLV slope between 5 and 20 Hz. In contrast to what was depicted with the detailed model, L5 and L6 PCs did not have high entrainment. However, the entrainment slope increased with tACS frequency, as shown in Figure 4.B. VIP displayed the highest PLV values and highest entrainment slope, especially for 5 and 10 Hz tACS. Beyond 10 Hz no increase was found. The lowest PLV variation was found for SST cells (Figure 4.B). Nonetheless, in the case of interneurons, the entrainment level was highly dependent upon the neocortical layer. Figure S1.1.B portrays the PLV variation for each interneuron type per layer, showing that almost all L23 interneurons have the highest entrainment slopes, which is consistent with lambda values provided in Table S.2.1

To visualize entrainment and especially the preferential phase at which activity was entrained, we computed phase histograms (displayed in Figure 5). In line with PLV results from the detailed model, pyramidal cells had the lowest intensity threshold for entrainment at the pseudo-endogenous stimulation frequency (10 Hz) where three of the five L5 PC were entrained at 1 V/m as well as three of the five L6 TPC (Rayleigh test, p¡0.05). At 2 V/m, all the L5 PC and four of the five L6 TPC

### Cell type specific response to tACS frequency

To further support this frequency dependency and validate this trend, using the detailed model, we performed the same simulation as in Figure 5, with varying stimulation frequency between 5-50 Hz in 5 Hz bins at an EF norm of 10 V/m (Figure 6). For most cells, we observed an entrainment at tACS frequency. While such high EF amplitude cannot be achieved using tACS devices for safety reasons [40], it enables drawing conclusions on frequency trends of tACS-induced neural entrainment. In line with previous results, L5 PC had maximal entrainment at 10 Hz, with the highest PLV values, while no clear maxima were observed for L6 TPCs and L23. However, the phase at which pyramidal cells were entrained was similar in trend for the three PC types, concavely increasing from around π/3 to 2π/3 (slightly before tACS waveform maxima to slightly after maxima). Interestingly, PV and SST cells had a mean increase in entrainment with increasing stimulation frequency, with higher mean entrainment for SST. The variability was higher than for excitatory cells, due to some unresponsive cells. The VIP cells had also a slight increase in entrainment, but were not as sensitive as other cells to tACS. Unlike PCs, the preferred phase was variable and had a non-monotone trend for the three considered inhibitory types over the considered spectrum.

**Figure 6:**
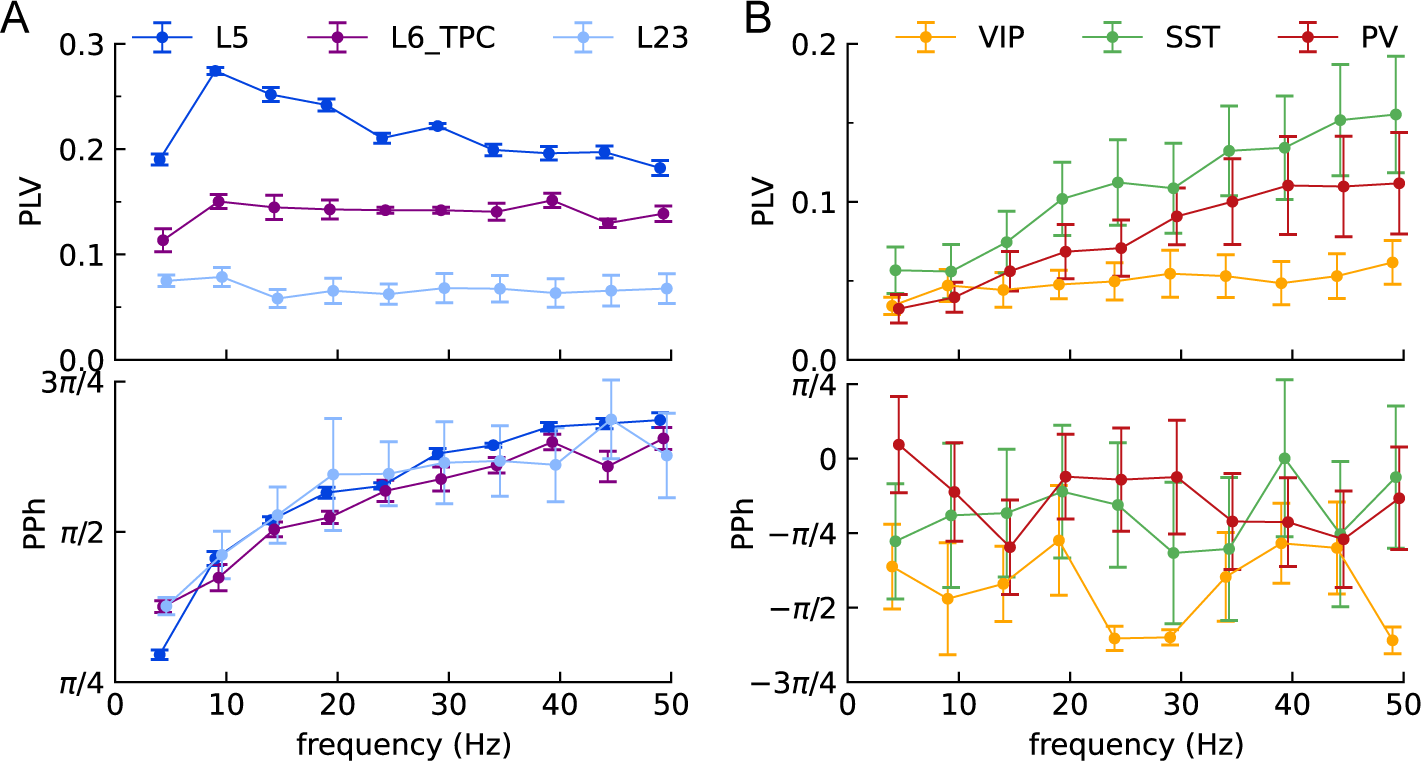
Spectrum of the PLV and associated preferred phase (PPh) for detailed excitatory cells (A) and for the three considered detailed inhibitory neuron types (B): VIP, SST, and PV. The plot shows the mean PLV and PPh with standard mean error (SEM) for each group.

To investigate further, we used the simplified model to map the spectrum of PLV and associated PPh for the frequency range (0-50 Hz, 2 Hz bins) with increasing intensity from 0 to 10 V/m. Figure 7 presents the quiver maps of the PLV and PPh variation for the main cell types. The Rayleigh test was used to delineate significant phase entrainments. For L2/3 PC, we can observe significant phase entrainment (PPh = 90°) from around 10 Hz for higher stimulation amplitudes (¿ 8 V/m), and then continue with increasing frequency for lower stimulation intensities (¿ 4 V/m). Throughout this tACS frequency and intensity ranges, the PPh deviated from 90° towards 45°, as depicted in Figure 6.A. For L5 and L6 PCs, we found similar behavior with a higher significant threshold for both frequency and intensity of stimulation. Moreover, the PPh of significant entrainment decreased from 270° to 235°. Regarding interneurons, SSTs had significant entrainment for 8-10 V/m amplitudes and higher PLVs for higher frequencies. PV and VIP cells had the highest PLV values around 10-20 Hz, with significant PLV values at lower EF intensities (2 V/m for VIP, and 4 V/m for PV).

**Figure 7:**
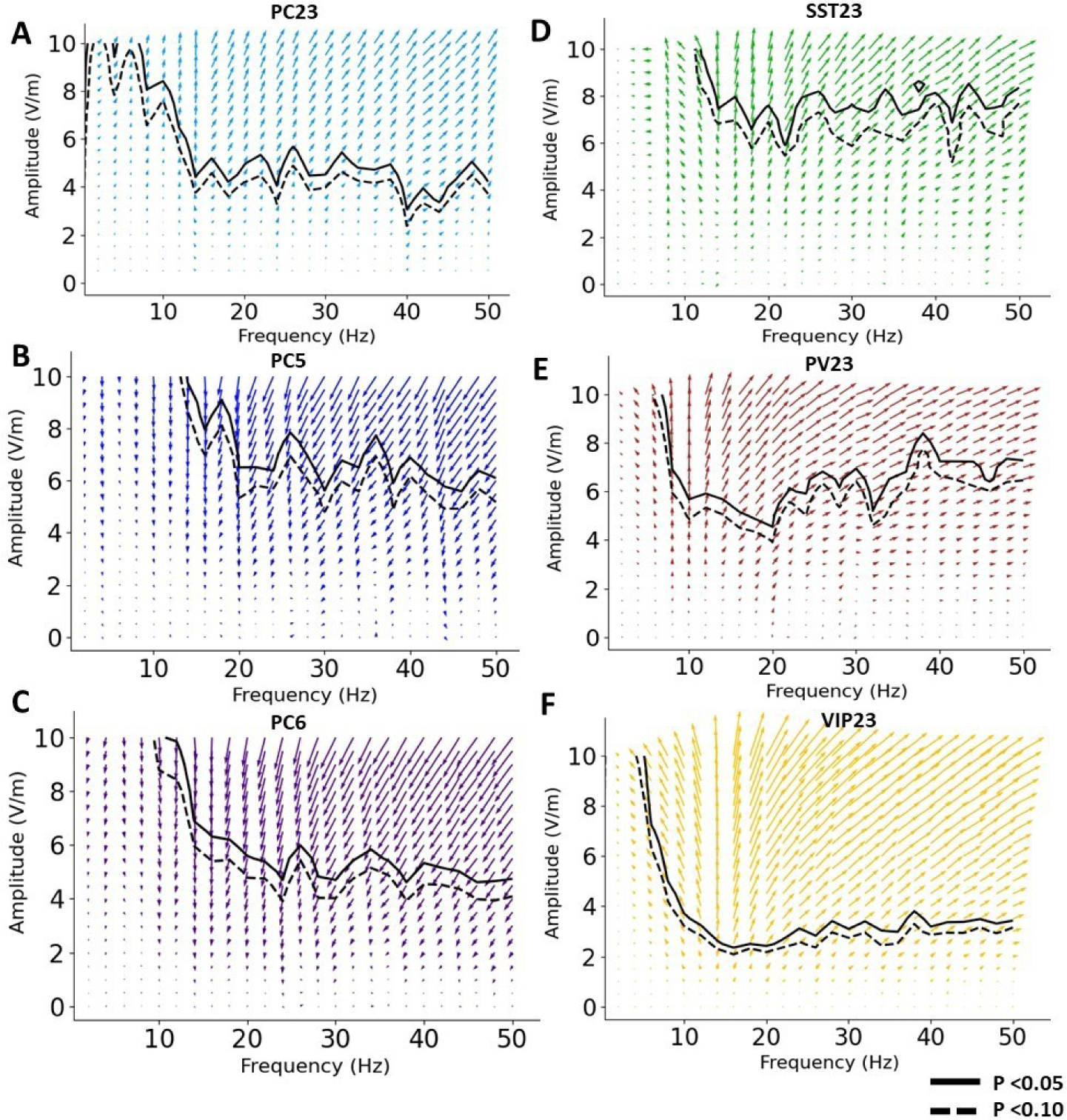
: Quiver plots of PLV variation with tACS frequency and amplitude for L23 (A), L5 (B), and L6 (C) PC cells and L23 PV (D), SST (E) and VIP (F) interneurons obtained using the simplified cell model. The arrow length and direction represent the Phase Locking Value (PLV) value and preferred phase (PPh), respectively. Significant phase entrainments were delineated for p ¡ 0.05 and p ¡ 0.01 using the Rayleigh test.

## DISCUSSION

To our knowledge, this is the first study to use two complementary types of realistic models to investigate the phase entrainment of neocortical cell types during tACS. This dual model approach serves two main purposes. First, it enabled studying cell-specific frequency responses to tACS, enhancing our understanding of tACS mechanisms and improving stimulation design for the specific stimulation of given cellular types. Second, it made it possible to compare results between detailed and simplified models. These simplified models are indeed valuable for faster computation and network modeling with properly calibrated EF coupling parameters, which could be used to study low-magnitude EF effects on an entire network. These models can also be useful in improving stimulation design at the level of a neural network, which is not possible with detailed models that include thousands of compartments.

Starting with the polarization spectrum computed for the detailed model, we found polarization length consistent with experimentally measured ones in [36], [37]. We observed that PCs demonstrate a type of low-pass filter behavior, akin to what was described in a prior study [12]. However, it has been reported that neurons may exhibit preferred frequencies, as experimentally demonstrated in hippocampal CA1 pyramidal cells [36]. These experiments revealed a similar exponential decay in the frequency of polarization constants, but with a peak occurring between direct current (DC) and 10 Hz, which differs from our findings. In contrast, the BP exhibited this frequency preference in the upper alpha band, possibly due to the increased conductance and contribution of delayed rectifier Kv3.1 channels [41]. PV and SST neurons exhibited a similar behavior, likely due to similar mechanisms. While some of them had a less distinct resonant frequency between 0 and 10 Hz, and a few did not exhibit resonant behavior at all, those cells generally demonstrated an increase in polarization length from DC, and maintained a nearly constant level across the remaining spectrum. To fine-tune the simplified models, which lack morphological detail, we used the polarization spectrum with the lambda values obtained from [34]. As an approximation, we assumed that each compartment of the simplified PC model had an identical polarization spectrum. Consequently, biophysics is crucial to explain the frequency dependency of EF sensitivity of cells.

Regarding phase entrainment, for the detailed model, L5 PCs had higher entrainment at their intrinsic firing rate frequency, further supporting the Arnold tongue mechanism [9], [19] since they were the most sensitive to EF for entrainment and polarization, with an effect threshold at 1 V/m for most the PC. Interestingly, this value of 1 V/m can be typically achieved in humans during tACS [42]. Nevertheless, it is interesting to note that close levels of entrainment between PC and inhibitory cells (SST and PV) were reached by increasing stimulation frequency. This supports the use of higher stimulation frequencies to selectively target inhibitory neurons. This result is in line with the results observed in [22], where the inhibitory network seems to be more sensitive to lower EF, while using higher EF switched inhibition to excitation, and therefore could have entrained pyramidal cells by reaching their entrainment threshold. These results further support the importance of biophysics when considering tACS since morphology did not necessarily induce higher entrainment. Additionally, VIP cells were not found to be reliably entrained at any frequency when modeled with synaptic inputs to generate a baseline activity around 10 Hz, while these cells had the highest polarization lengths of the tested inhibitory cells. This can be explained by the high synaptic weight coefficient required to produce a 10 Hz firing rate, which provided a strong input compared to the contribution of the external weak electric field (see synaptic weight coefficients of Table S.2.1). Therefore, entrainment is input-dependent, which is consistent with recent reported results in non-human primates [21], where strongly entrained neurons were less entrained to tACS. This highlights the need for validation in physiological activity conditions, with maybe an even more realistic set of synapses and pre-synaptic activity. Also, this suggests that findings from studies using external sources to generate neural activity must be taken with care to draw conclusions on tACS entrainment pattern since, for example, monophasic inputs could be easier to bias with a small alternating input than random events. Although PLV is known to be a biased estimate [43], it remains a valuable metric used in other related studies [16], [17], [24]. Therefore, the choice of PLV in the present study enabled a direct comparison of results.

It is important to highlight that both detailed and simplified models exhibit similar responses to the electric field. In both model types, PLV increased linearly with electric field amplitude. However, it appears that the EF applied to the L23 pyramidal cell had a more pronounced effect on the simplified model as compared to the detailed model. This difference may originate from the lambda values that we used, which are the highest for the dendritic compartment of the L23 pyramidal cell. The simplified model enabled mapping the 2D PLV variation with EF intensity and frequency for the considered cells. As depicted in Figure 7, the quiver plots revealed the ranges in which each cell type’s spiking activity is significantly entrained to the tACS-induced EF. PCs mainly exhibited strong spike-field entrainment to EF that increased with frequency at alpha and beta bands, and was stable for higher frequencies. This response to tACS required a lower EF intensity threshold for L23 PCs than for L5 and L6 PCs. In contrast, quiver plots of inhibitory neurons displayed phase entrainment that was the highest between 10 and 30 Hz, and then decreased at higher frequencies. Moreover, VIP cells demonstrated the highest sensitivity to the weak EF, starting around a threshold of 2 V/m. Finally, we have to mention that for all cell types, the PPh is dependent on the frequency of stimulation (Figure 7).

These results highlight how tACS interacts with different cell types. However, it is worth noting that cells were not coupled in this study. Indeed, here we investigated the cell-specific responses to tACS that could explain their potential role in network effects. Furthermore, the range of tACS amplitude explored was higher than EF achieved in real-life experiments but attempting at identifying a threshold for an effect on all cells. It was previously shown that isolated cells have higher thresholds for EF effect than when they were coupled [44], implying that the threshold values identified in our study are probably higher than if cells would have been coupled through synaptic projections for example Also, ephaptic interactions were not included but should be considered to accurately predict network effects [11], [14]. Therefore, the reported threshold here likely reflects a worst-case scenario. However, the threshold for an effect was observed in the 1-2 V/m, which is actually close to what could be achieved during *in-vivo* experiments in rodents and primates [42]. Further work would consider coupled simplified cells, since they were shown to properly incorporate realistic responses to tACS and benefit from low computational cost compared to realistic cells. Another potential direction would be to incorporate synaptic terminal polarization and further consider the synaptic vesicle release probability change induced by tACS, as it has been done for tDCS [45], which could further lower the threshold. BBP-generated cells were used for realistic modeling, but the axon was cut prior to fitting, and therefore the polarization length could be different from the original cell. For instance, [34] reported other polarization lengths using the full axonal arbor tree of the same set of cells, but conductance was not optimized on the full tree, which can introduce a bias. Additionally, it might be valuable to evaluate an axonal tree incorporating a myelin mechanism, similar to previous efforts [25]. However, to our knowledge, no precise model involving realistic cells together with myelin mechanism could be used to assess the change to EF sensitivity, since non-causal behavior was found using the reported model in [24], [25] with little to no EF (see Figure S2.11). Therefore, efforts are needed to build a precise model with the full axonal tree fitted to experimental measures, with and/or without myelin. Conducting such an analysis on a realistic model that incorporates mechanisms specifically attributed to human neocortex cells would also be of interest, especially with backpropagation mechanisms and calcium channels distributed on the dendritic tree.

## CONCLUSIONS

Using both detailed and simplified neural models with a variety of inhibitory neurons, we demonstrated the ability of higher tACS frequencies to recruit specific classes of inhibitory cells, while pyramidal cells were more sensitive to their own firing rate frequencies in the detailed models, supporting the Arnold tongue phenomenon. Simplified and realistic models were in relatively good agreement in terms of entrainment threshold, opening the possibility to use simplified models to study further network effects of tACS with realistic effects at the single cell level using coupling constants based on realistic neuronal morphologies.

## ACKNOWLEDGMENTS

This work has received a French government support granted to the CominLabs excellence laboratory and managed by the National Research Agency in the ”Investing for the Future” program under reference ANR-10-LABX-07-01; and from the European Research Council (ERC) under the European Union’s Horizon 2020 research and innovation programme (grant agreement No 855109).

## S.1: THE SIMPLIFIED SINGLE-CELL MODELS’ EQUATIONS, PARAMETERS, AND RESULTS

As previously mentioned, the three-compartments neuron model is based on the work of [1]. In this new version, we added a third compartment that portrays the AIS wherein the corresponding voltage-dependent ionic currents consisted of the sodium (i*_Na_*) and the potassium (i*_K_*) currents in addition to the leak current. The corresponding equations were adapted from (Traub et al. 2003). The coupling of the soma/dendrites compartments and Soma/AIS was made through the conductances g*_SD_* and g*_SA_*, respectively (Figure 2.A). Ionic channel parameters’ values were fine-tuned to replicate the firing rate observed in electrophysiological recordings. A comprehensive explanation of the equations governing ionic currents and their associated parameter values can be found in [1]. The values for the conductance and reversal potentials are given in Tables A.1.1 and A.1.2.

The membrane potentials of the somatic (V*_s_*), dendritic (V*_d_*) and AIS (V*_A_*) compartments were depicted as follows:

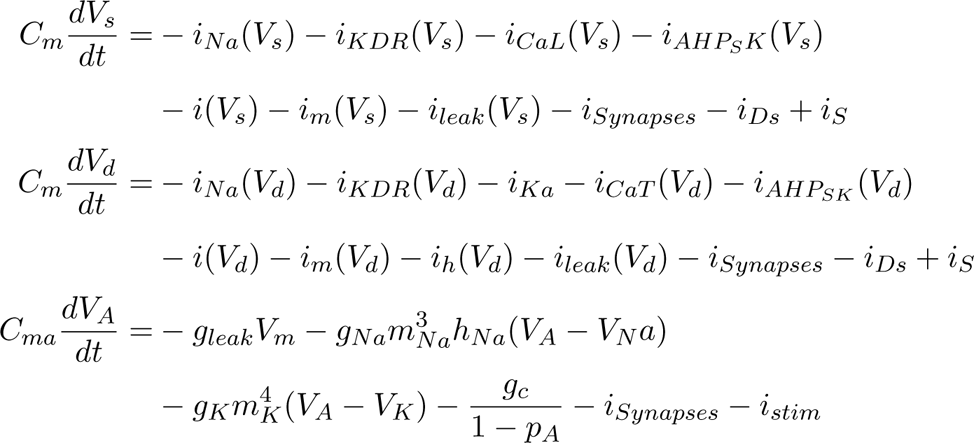

With i*_N_*a the voltage-dependent sodium current, i*_K_* the potassium current, i*_KDR_* the potassium delayed-rectifier current, i*_AHP_* the calcium-dependent potassium currents, i*_m_* the muscarinic current, i*_CaL_*, i*_CaR_* and i*_CaT_*the L/R/T-type calcium currents, i*_KA_* the fast inactivating potassium current, i*_h_* the hyperpolarization-activated cationic current, i*_leak_* is the leak current, i*_KNa_* the sodium-activated potassium current, i*_KCa_* the calcium-activated potassium current, i*_D_* and i*_s_*are the injected input currents of the dendritic and somatic compartments respectively, i*_Synapses_*: are the excitatory and inhibitory postsynaptic currents, i*_SD_* is the current entering the dendrites from the soma 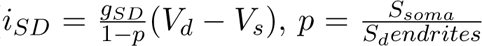 the ratio between the surface of the soma and dendrites, p = 0.15 ), i*_DS_* is the current entering the soma from the dendrites 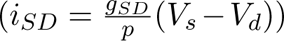).

**Table S.1.1:** Values for the ionic currents conductances for the cells modeled using the simplified reduced single-cell model.

| Compartment |  | Conductance mS/cm2 |  |  |  |  |
| --- | --- | --- | --- | --- | --- | --- |
|  |  | Layer 2/3 | Layer 5 | PV<sub>+</sub> | SST+ | VIP+ |
| Dendrites | $i_{Na}$ | 14 | 14 | - | - | - |
| | $i_{KDR}$ | 2 | 2 | - | - | - |
| | $i_{CaT}$ | 1 | 1 | - | - | - |
| | $i_{CaR}$ | 3 | 0.75 | - | - | - |
| | $i_{AHP}$ | 10 | 20 | - | - | - |
| | $i_{BK}$ | 1 | 1 | - | - | - |
| | $i_m$ | 0.1 | 0.1 | - | - | - |
| | $i_h$ | 0.4 | 0.4 | - | - | - |
| | $i_{KA}$ | 55 | 55 | - | - | - |
| | $i_K$ | - | - | - | - | - |
| Soma | $i_{Na}$ | 70 | 70 | 35 | 35 | 35 |
| | $i_{KDR}$ | 6 | 6 | 9 | 9 | 9 |
| | $i_{CaL}$ | 0.5 | 0.5 | - | 1 | - |
| | $i_{AHP}$ | 0.1 | 0.1 | - | - | - |
| | $i_{BK}$ | 2 | 2 | - | - | - |
| | $i_m$ | 3.1 | 3.5 | - | - | - |
| | $i_{KNa}$ | 0 | 0.5 | - | - | 1.33 |
| | $i_h$ | - | - | - | 0.15 | - |
| | $i_{KCa}$ | - | - | - | 10 | - |
| AIS | $i_{Na}$ | 200 | 200 | - | - | - |
| | $i_{KDR}$ | 200 | 200 | - | - | - |

**Figure S.1.1:**
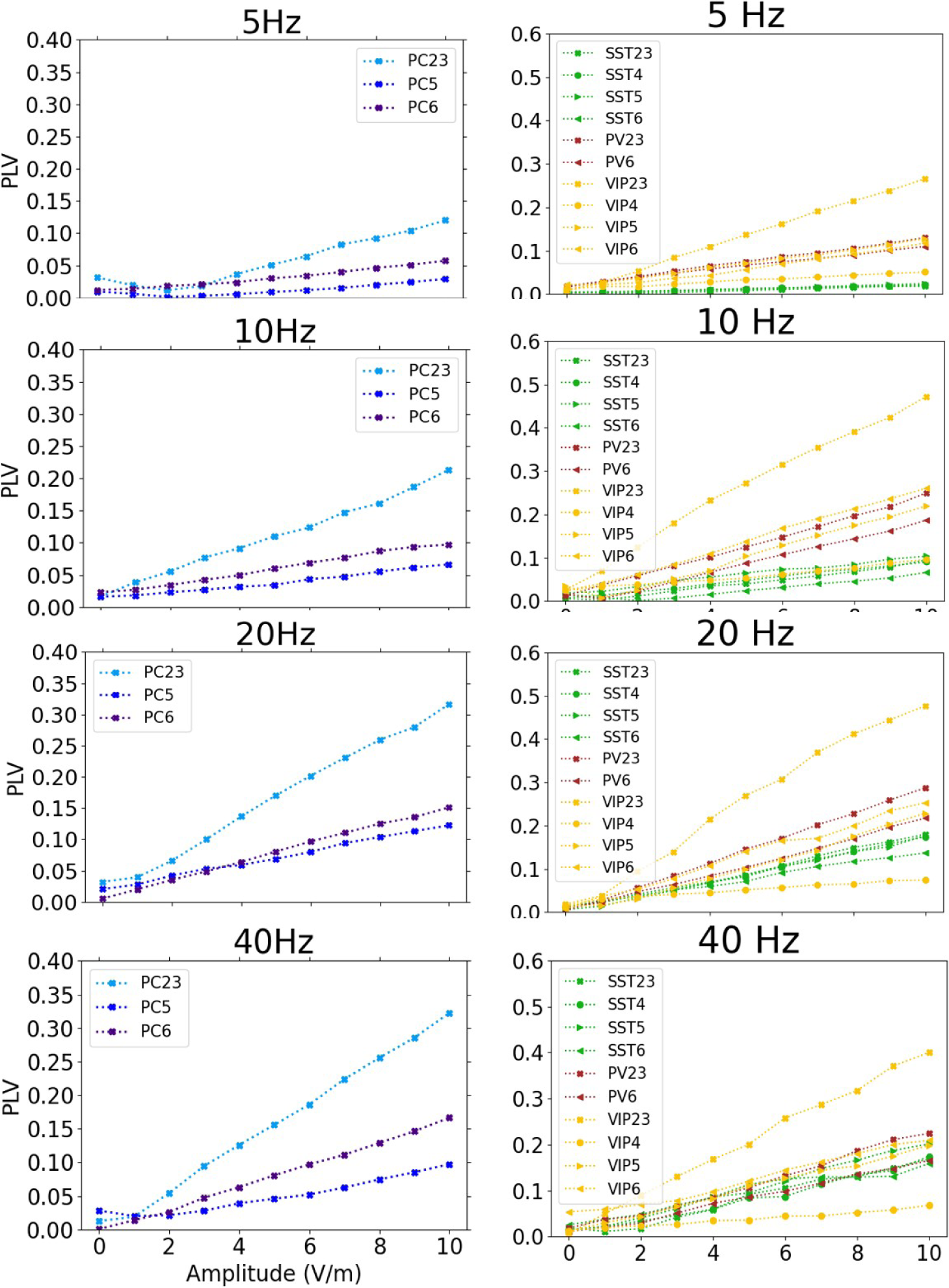
Individual PLV dose-response curves for PC L2/3, L5, L6 cells (left) and interneurons VIP, SST, and PV L2/3, L5, L6 for 5, 10, 20 and 40 Hz–tACS.

**Table S.1.2:**
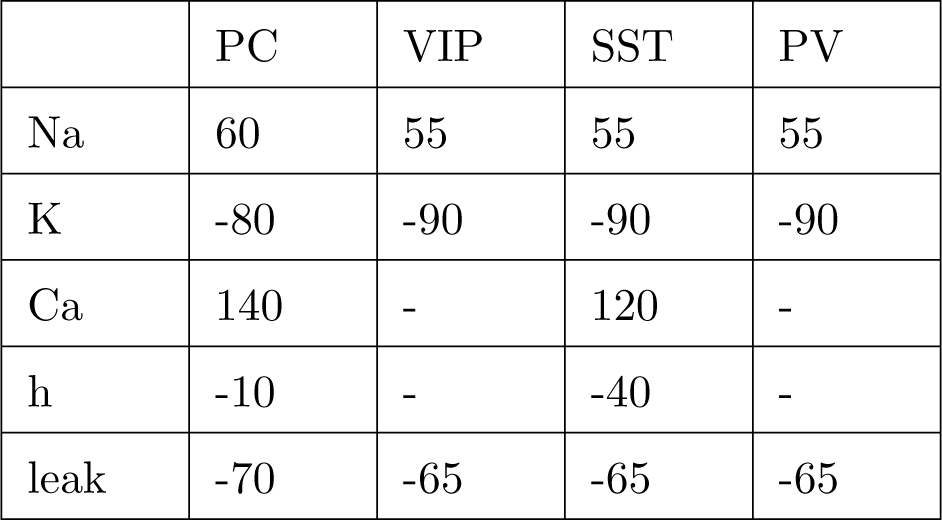
Values for the reversal potentials of voltage-gated channels of the cells modeled using the reduced cell model.

|  | PC | VIP | SST | PV |
| --- | --- | --- | --- | --- |
| Na | 60 | 55 | 55 | 55 |
| K | -80 | -90 | -90 | -90 |
| Ca | 140 | - | 120 | - |
| h | -10 | - | -40 | - |
| leak | -70 | -65 | -65 | -65 |

## S.2: THE DETAILED SINGLE-CELL MODELS’ CHARACTERISTICS AND SIMULATION RESULTS

**Table S.2.1:**
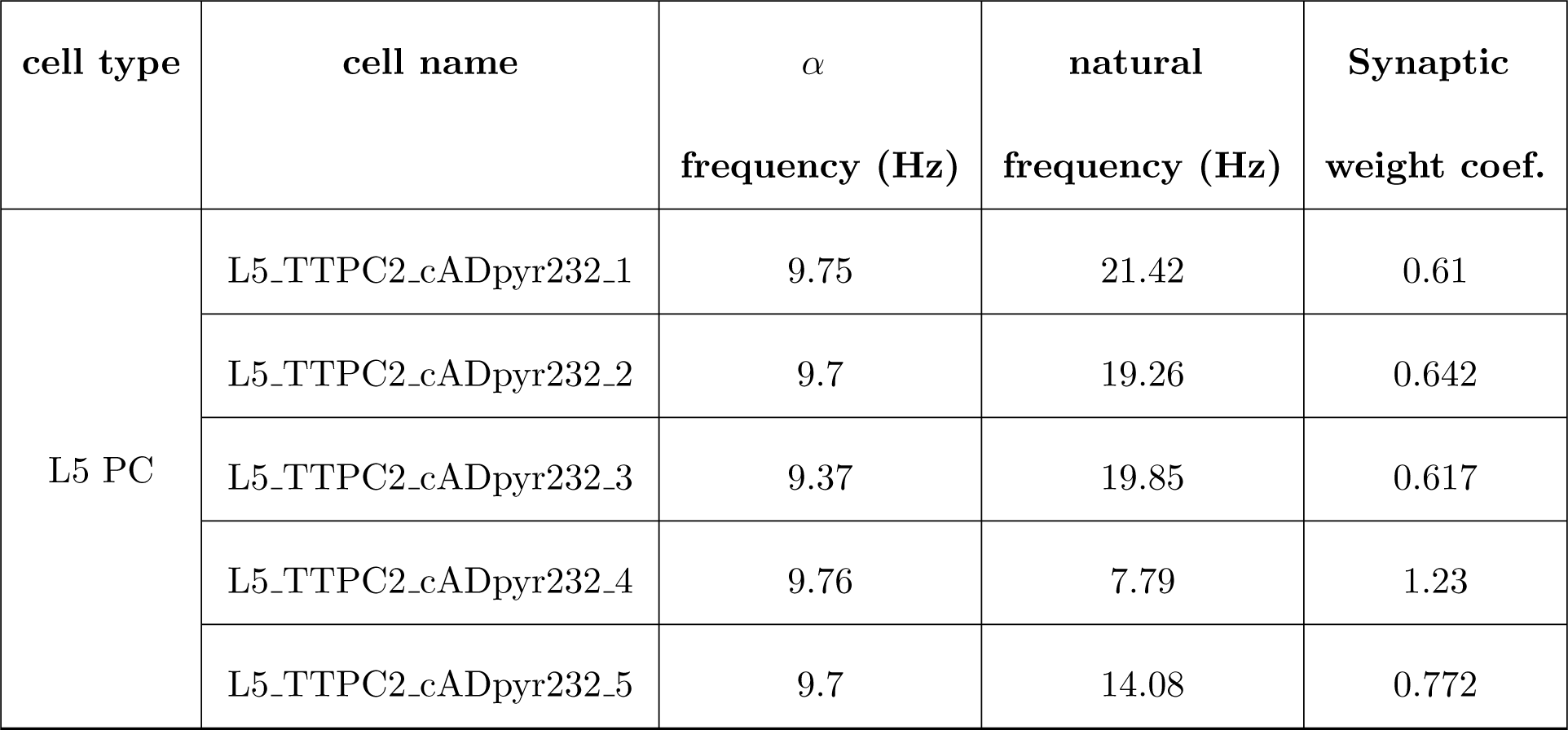
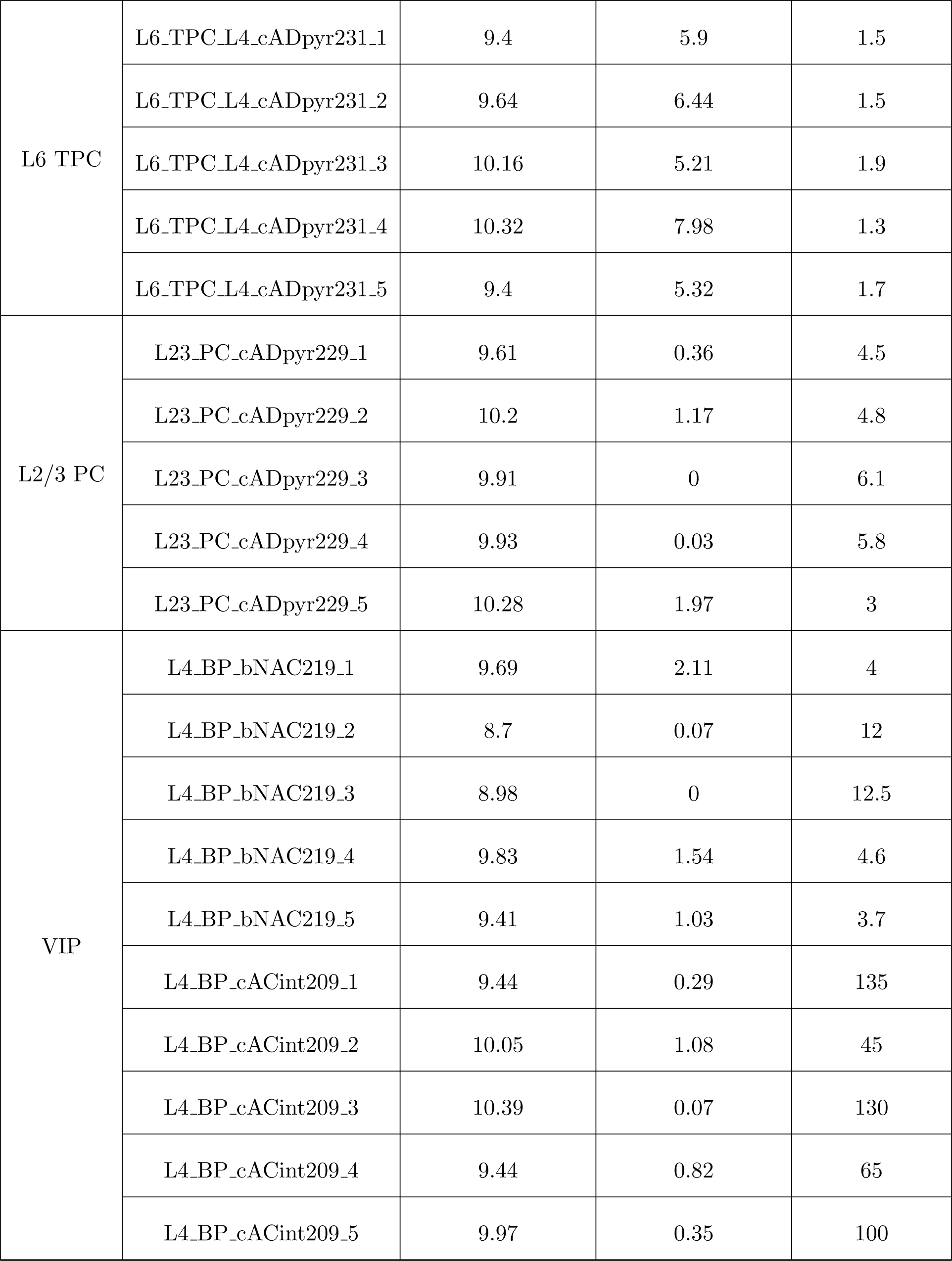
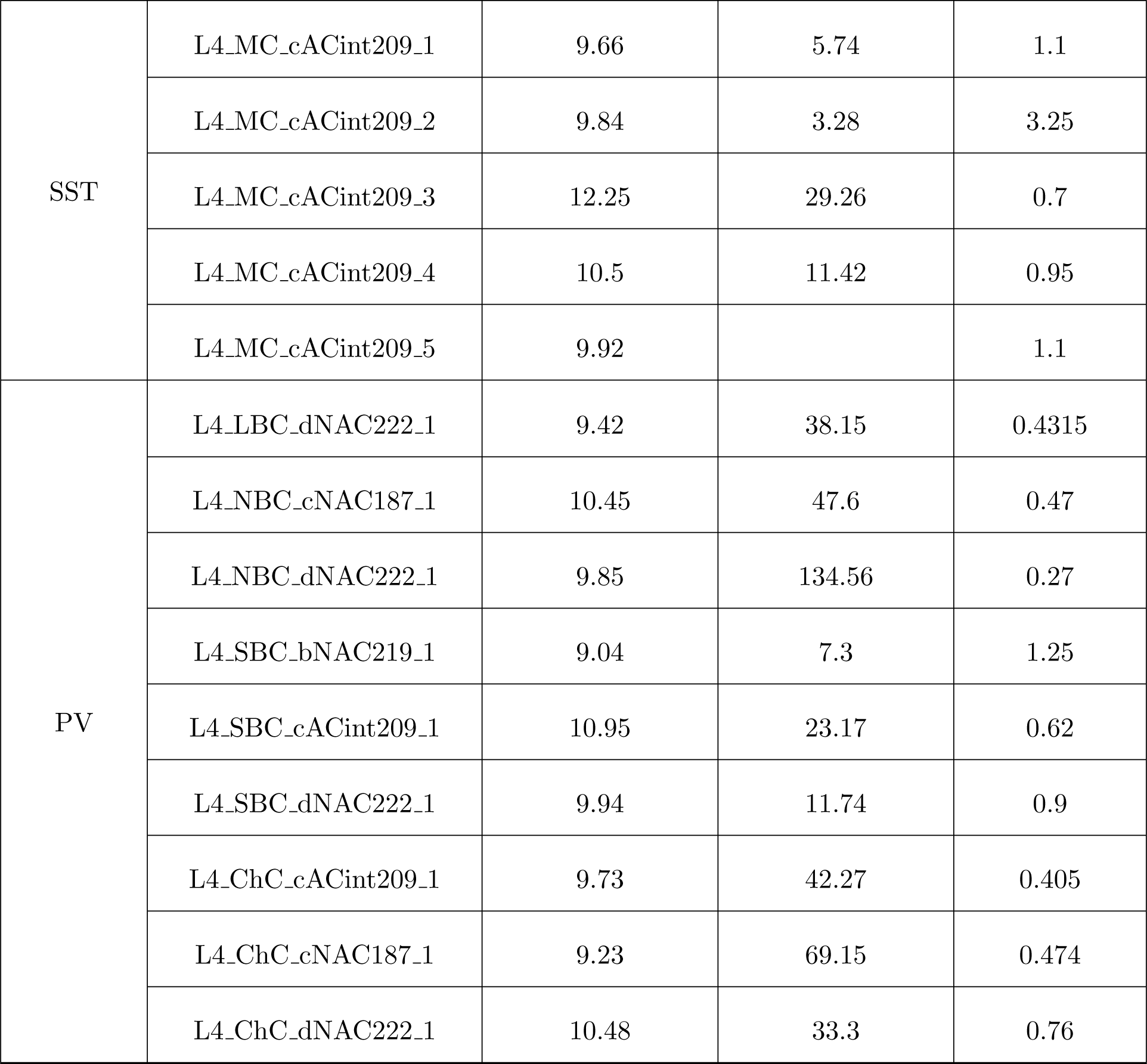
Cell type considered and their associated name from BBP cells. The baseline frequency with the synaptic input to drive 10 Hz activity is provided, as well as the frequency with BBP initial synaptic weights.

**Figure S.2.1:**
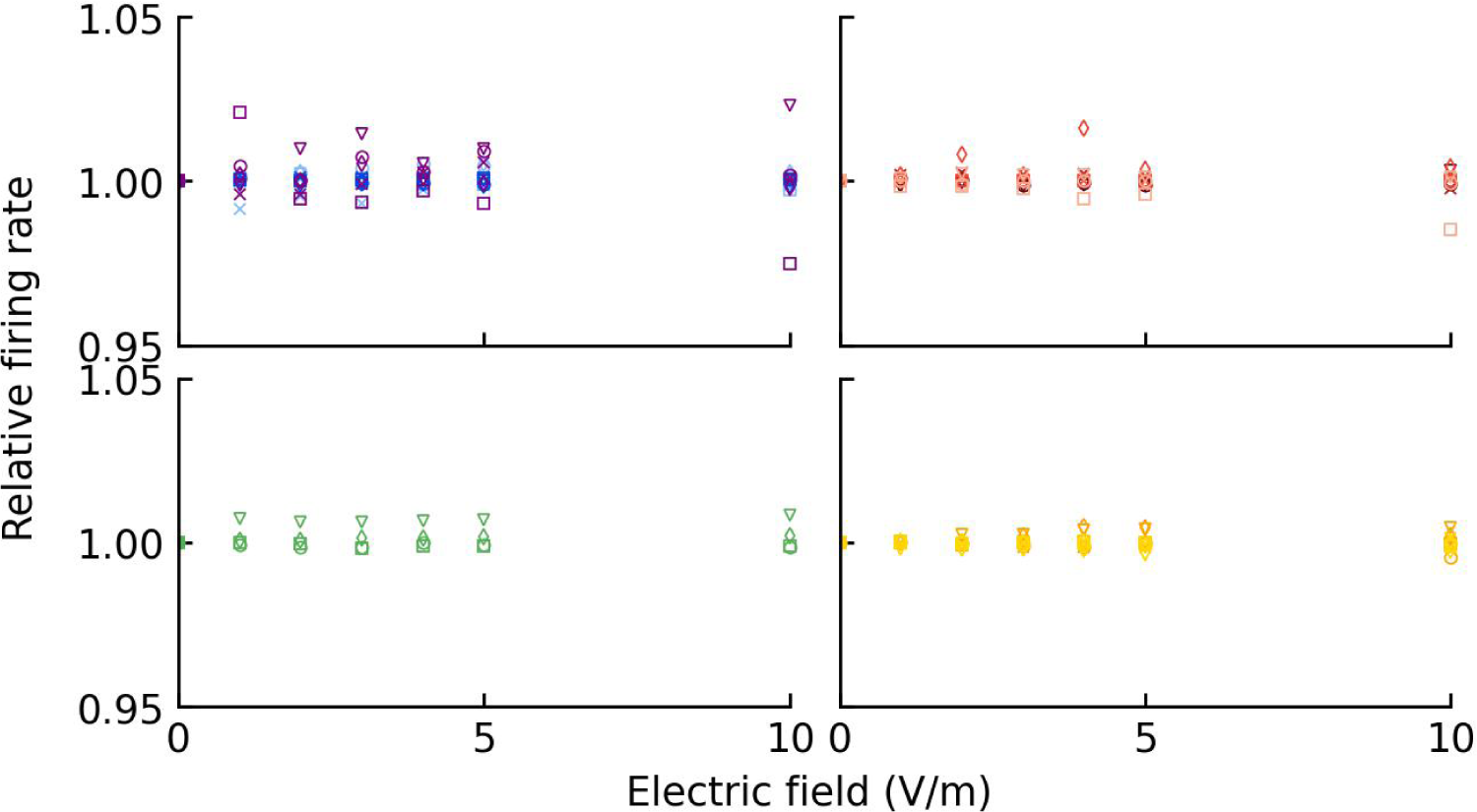
Relative firing rate plot during 5 Hz-tACS with different amplitudes. No clear effect on FR can be observed for all cellular types.

**Figure S.2.2:**
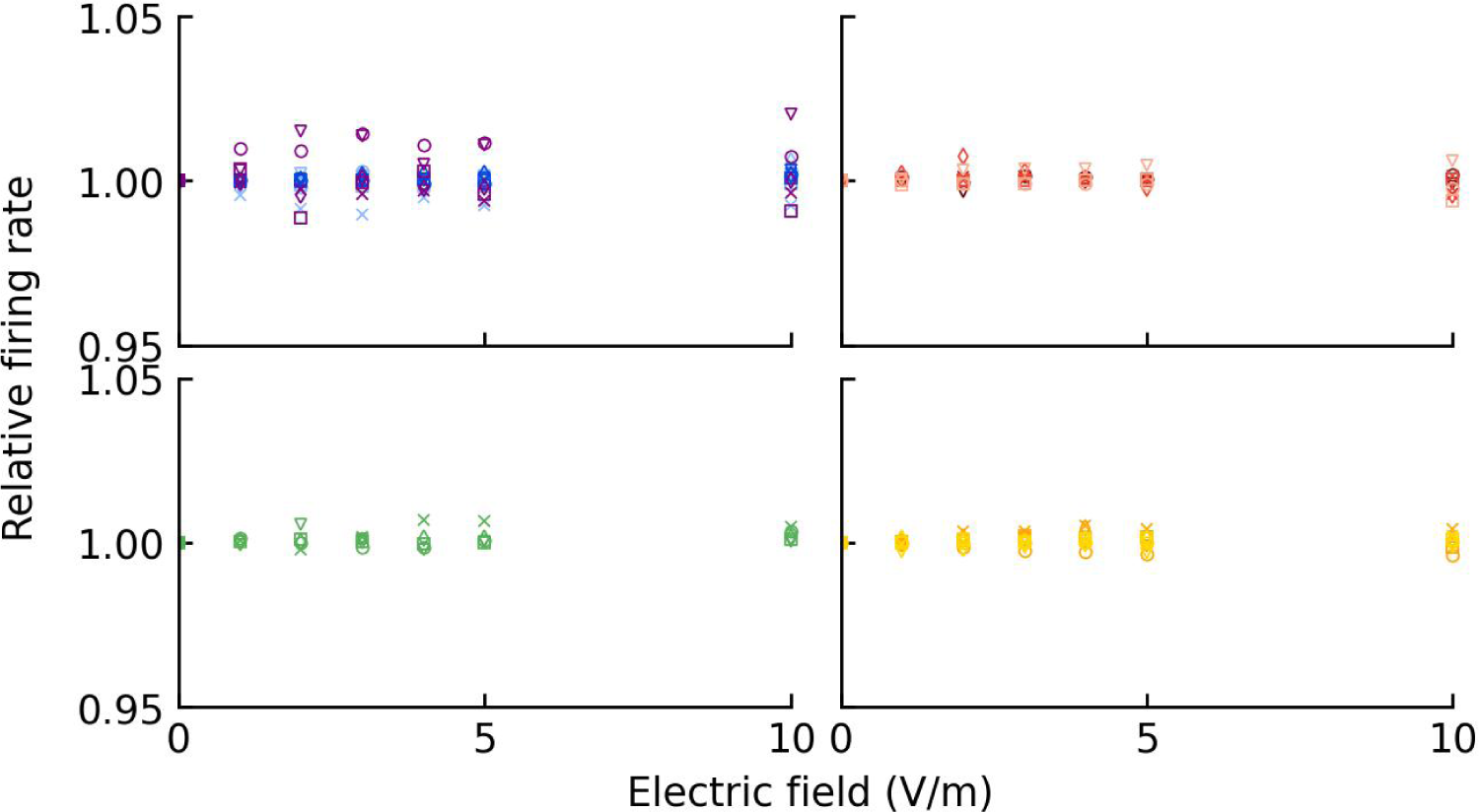
Relative firing rate plot during 10 Hz–tACS with different amplitudes. No clear effect on FR can be observed for all cellular types.

**Figure S.2.3:**
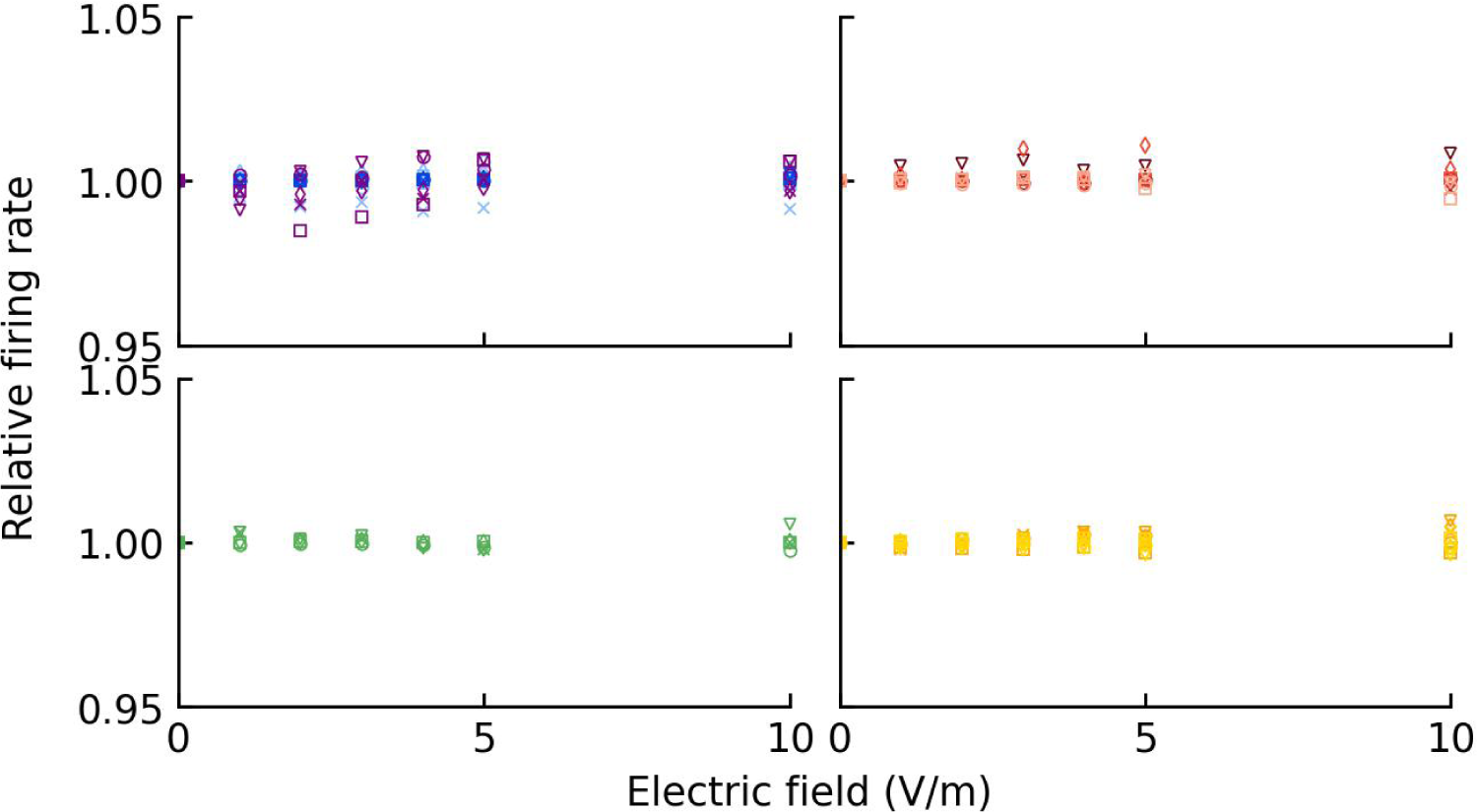
Relative firing rate plot during 20 Hz–tACS with different amplitudes. No clear effect on FR can be observed for all cellular types.

**Figure S.2.4:**
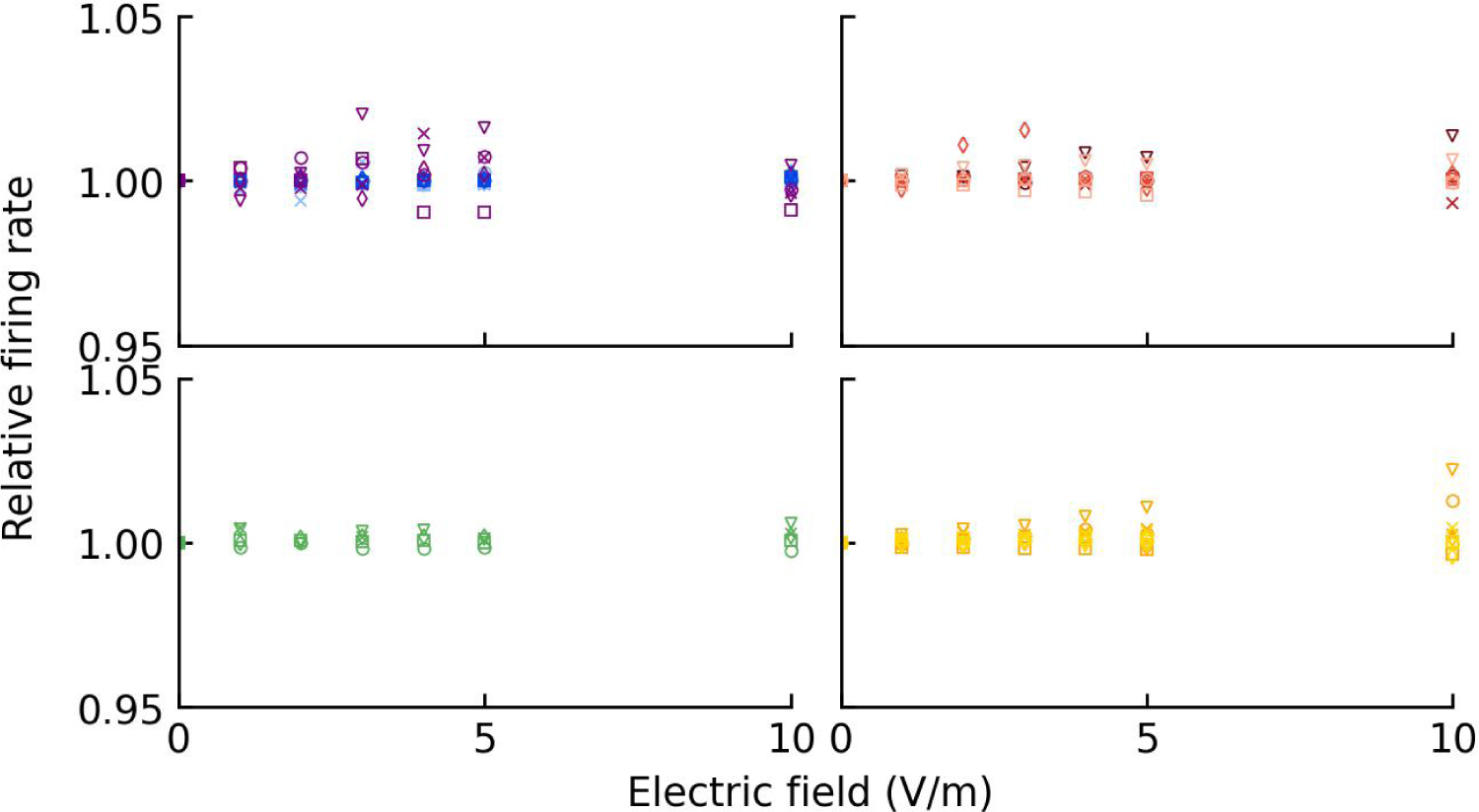
Relative firing rate plot during 40 Hz–tACS with different amplitudes. No clear effect on FR can be observed for all cellular types.

**Figure S.2.5:**
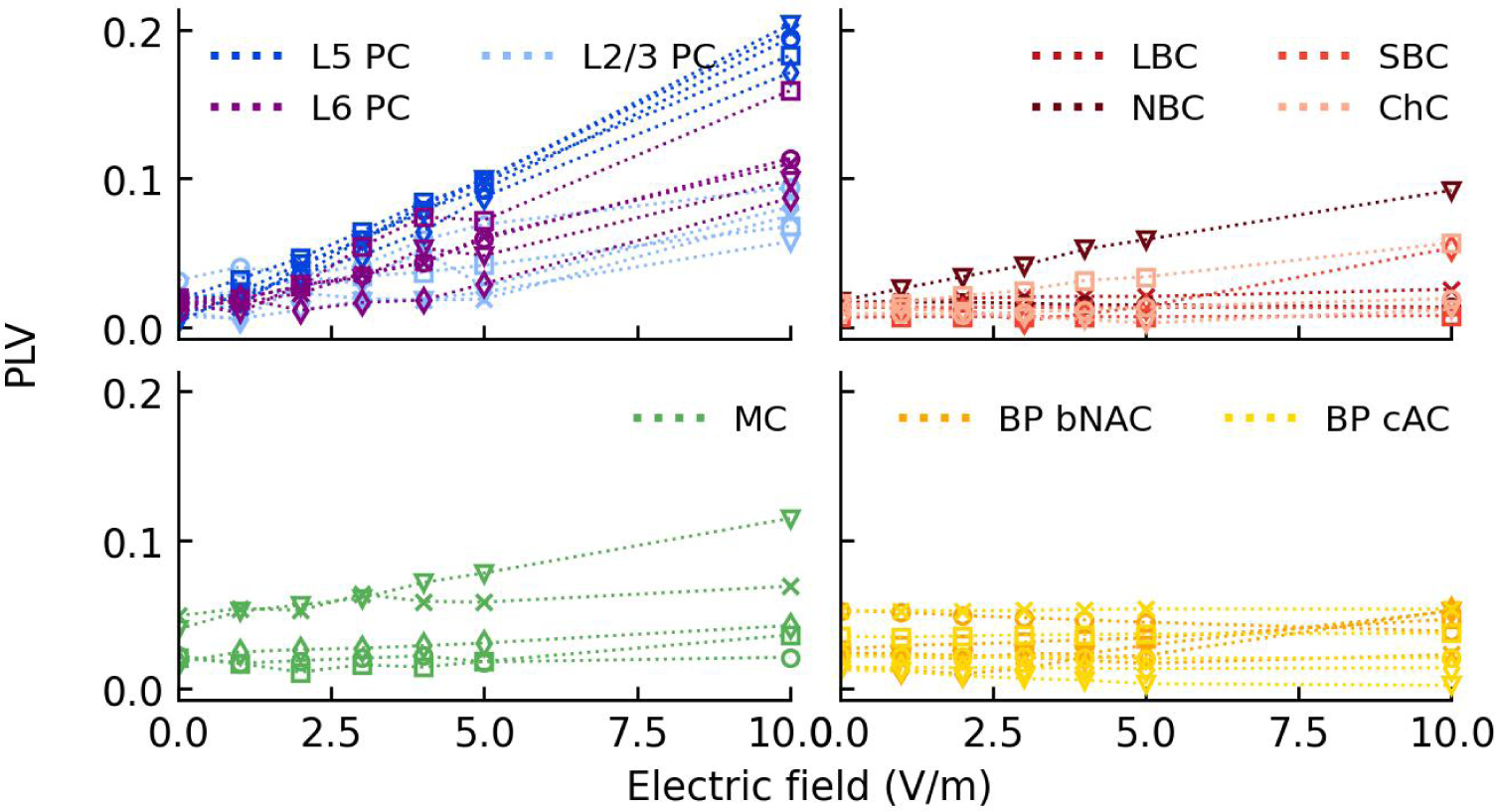
Individual PLV dose-response curves for PC (L2/3, L5, L6), VIP, SST, and PV for 5 Hz–tACS.

**Figure S.2.6:**
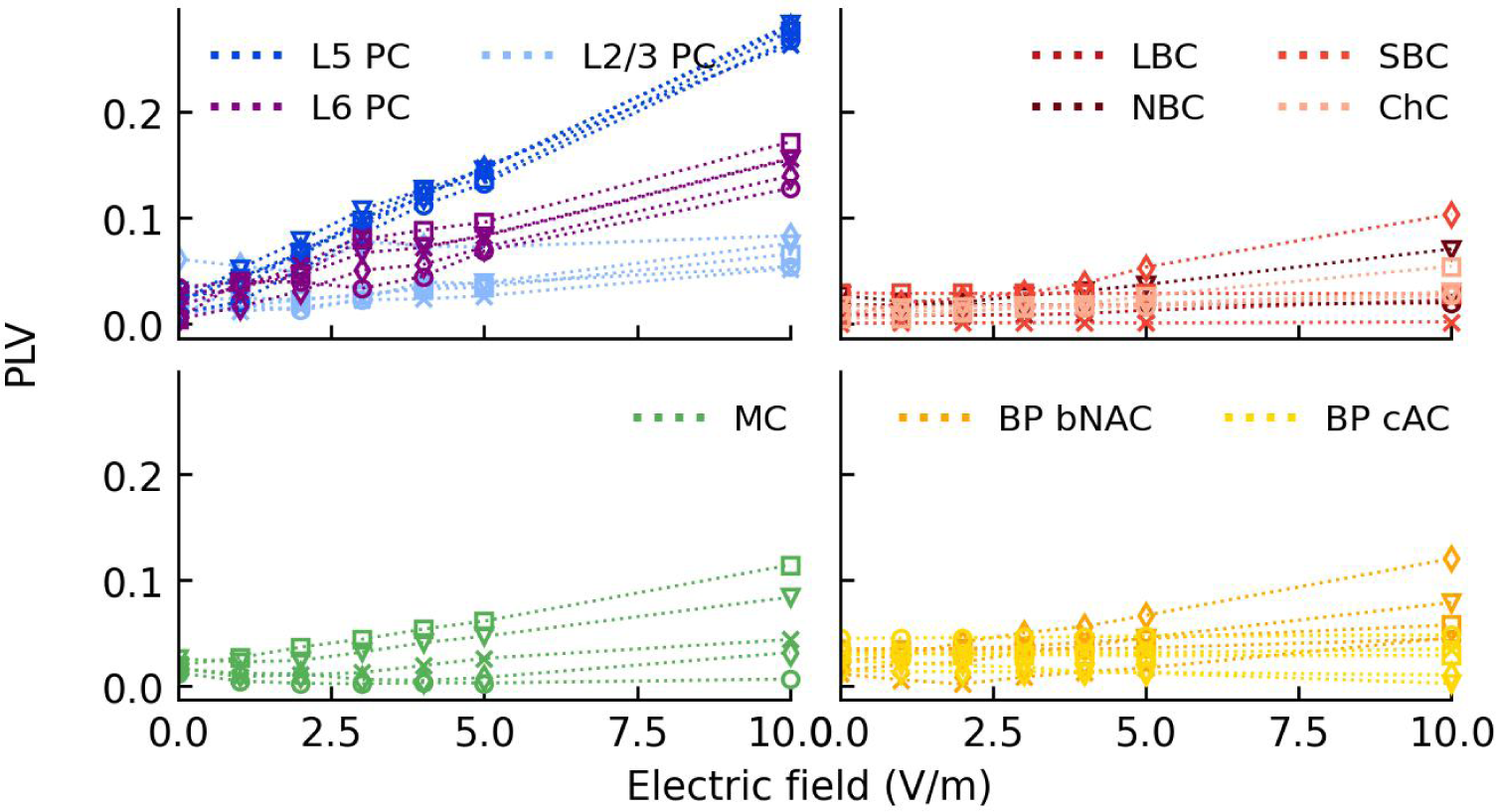
Individual PLV dose-response curves for PC (L2/3, L5, L6), VIP, SST, and PV for 10 Hz–tACS.

**Figure S.2.7:**
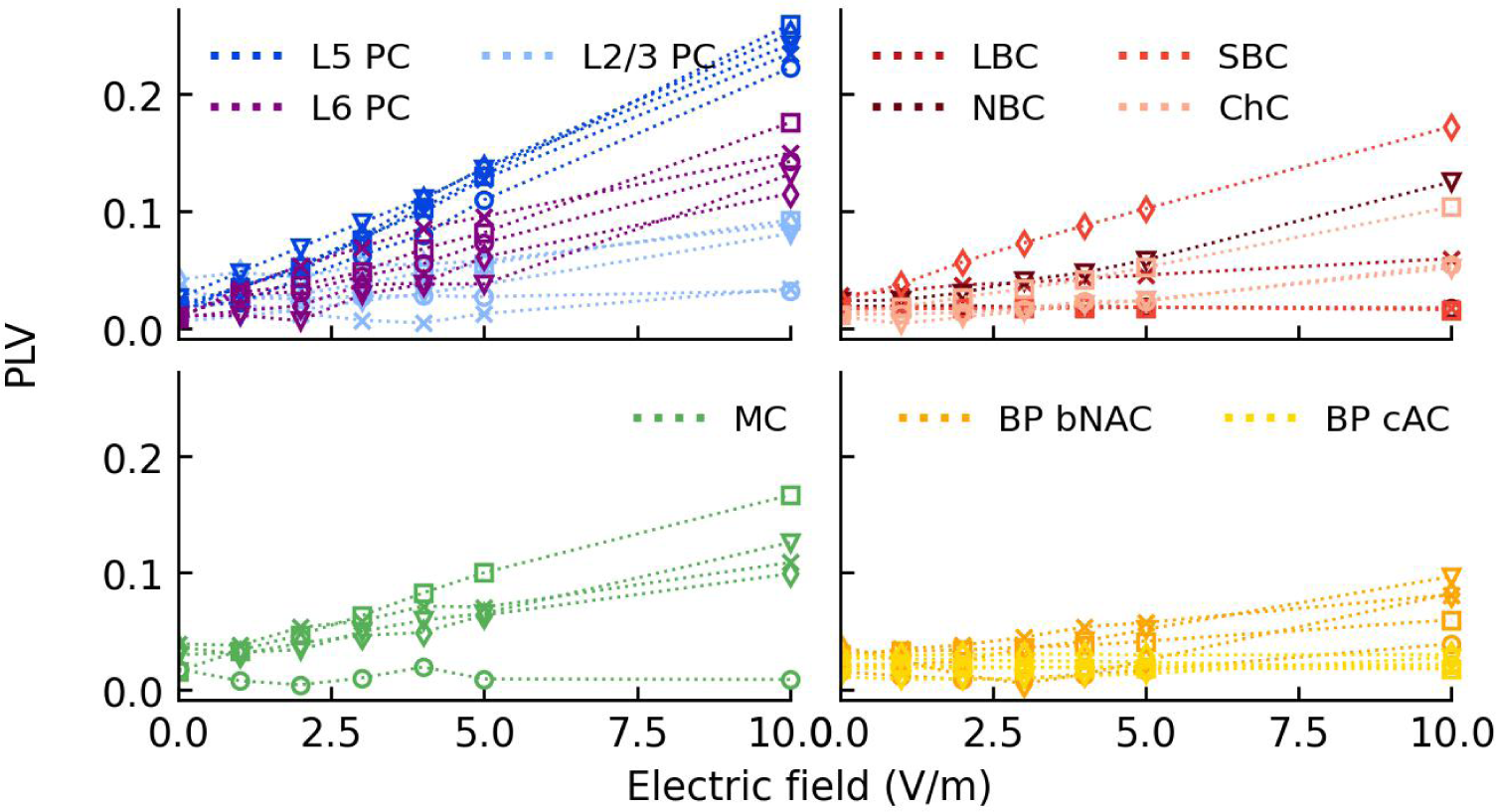
Individual PLV dose-response curves for PC (L2/3, L5, L6), VIP, SST, and PV for 20 Hz–tACS.

**Figure S.2.8:**
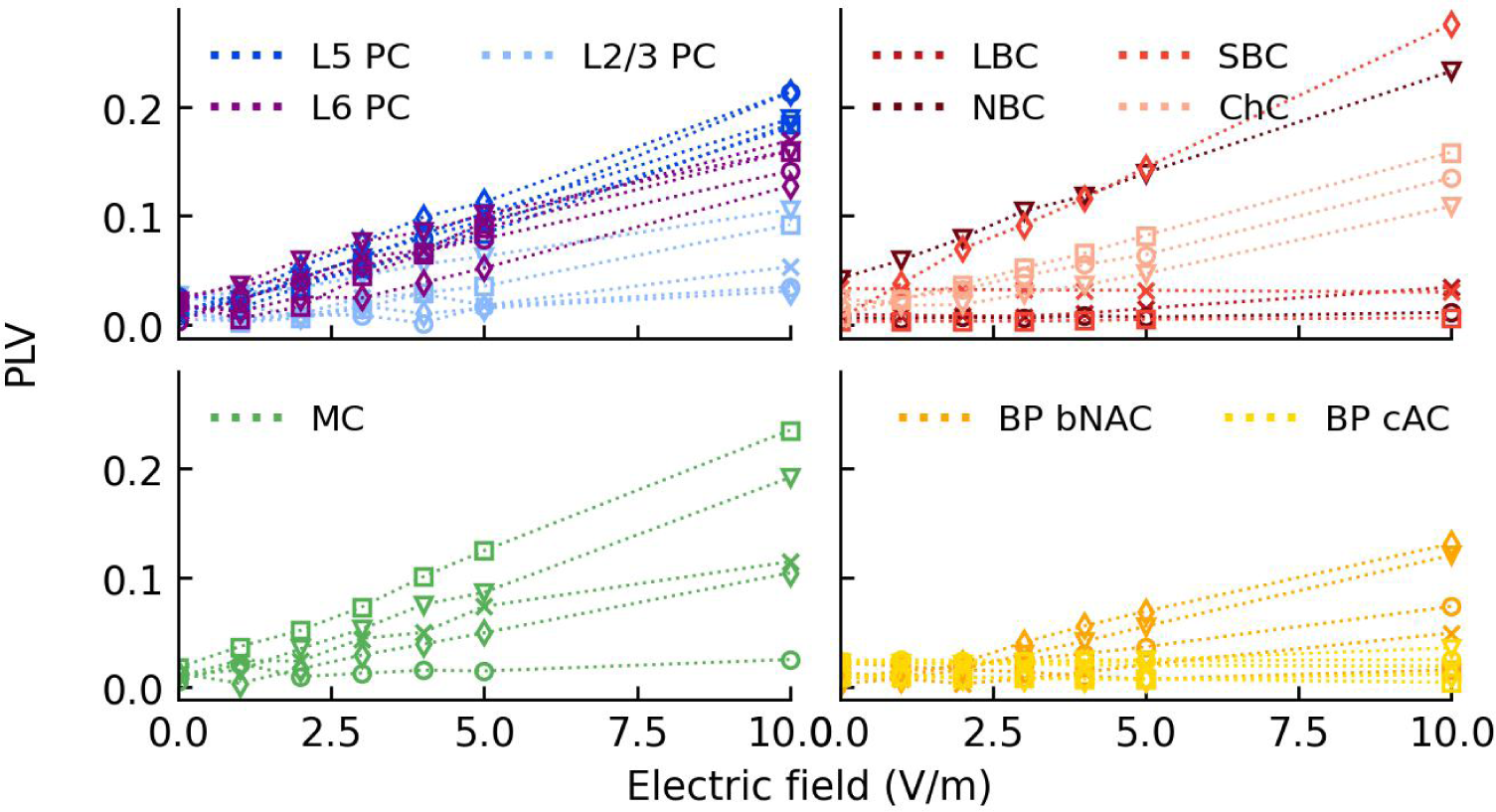
Individual PLV dose-response curves for PC (L2/3, L5, L6), VIP, SST, and PV for 40 Hz–tACS.

**Figure S.2.9:**
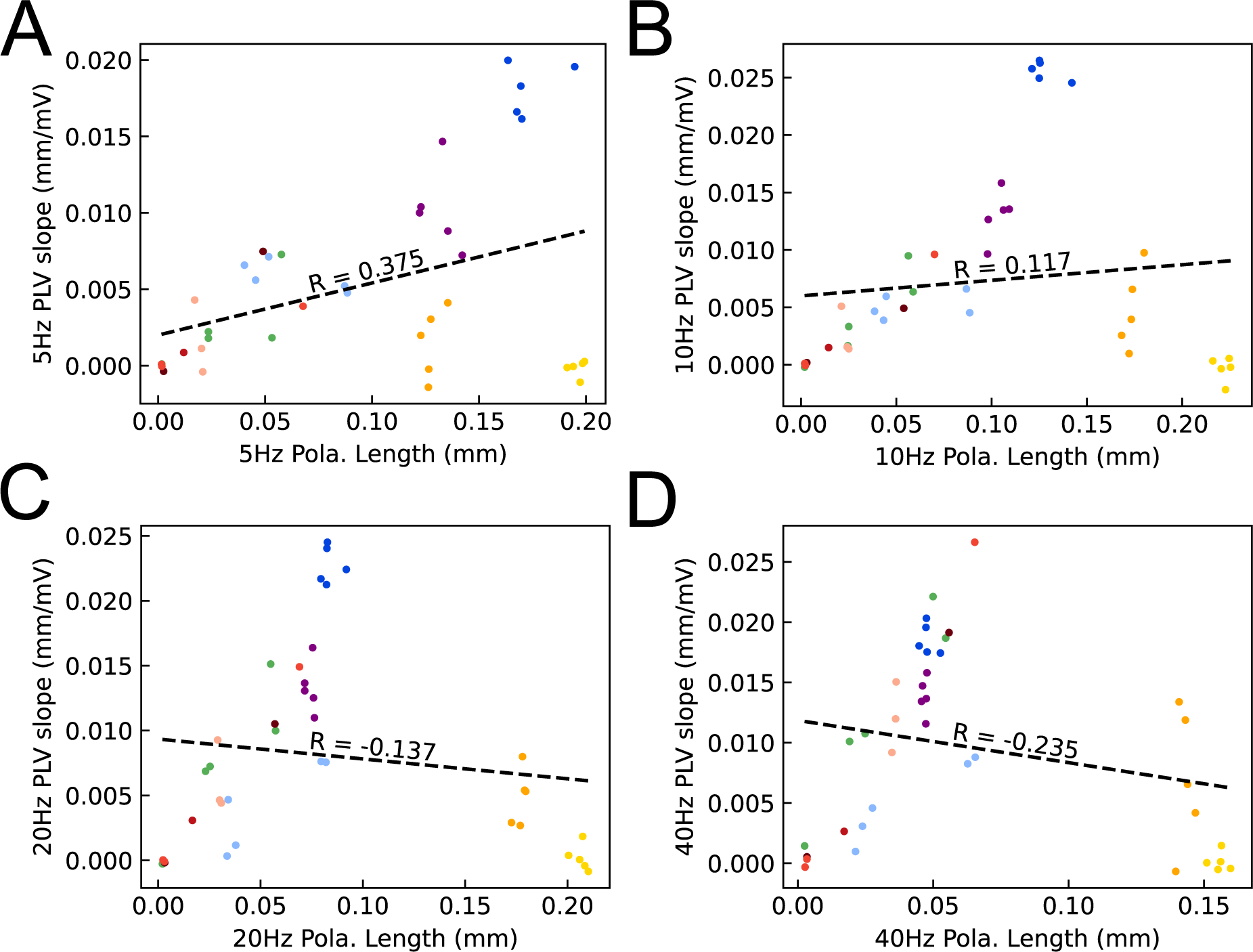
Relations between PLV slope and polarization length for 5 (A), 10 (B), 20 (C), and 40 Hz (D) stimulation frequency with 10 Hz baseline activity. Pearson’s coefficients are indicated on the plots which were all significant (p¡1e-3).

**Figure S.2.10:**
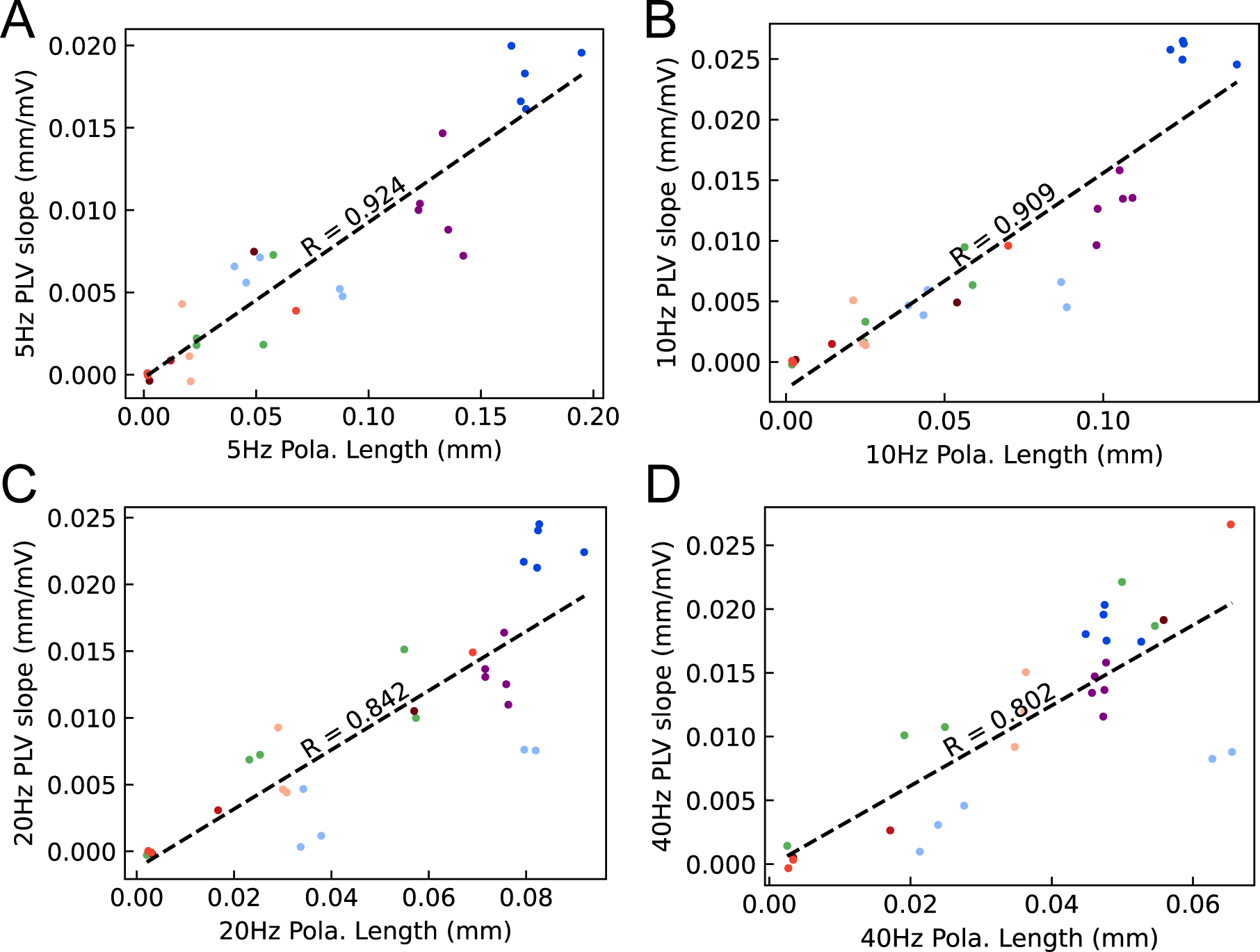
Relations between PLV slope and polarization length excluding VIP cells for 5 (A), 10 (B), 20 (C), and 40 Hz (D) stimulation frequency with 10 Hz baseline activity. Pearson’s coefficients are indicated on the plots which were all significant (p¡1e-3).

**Figure S.2.11:**
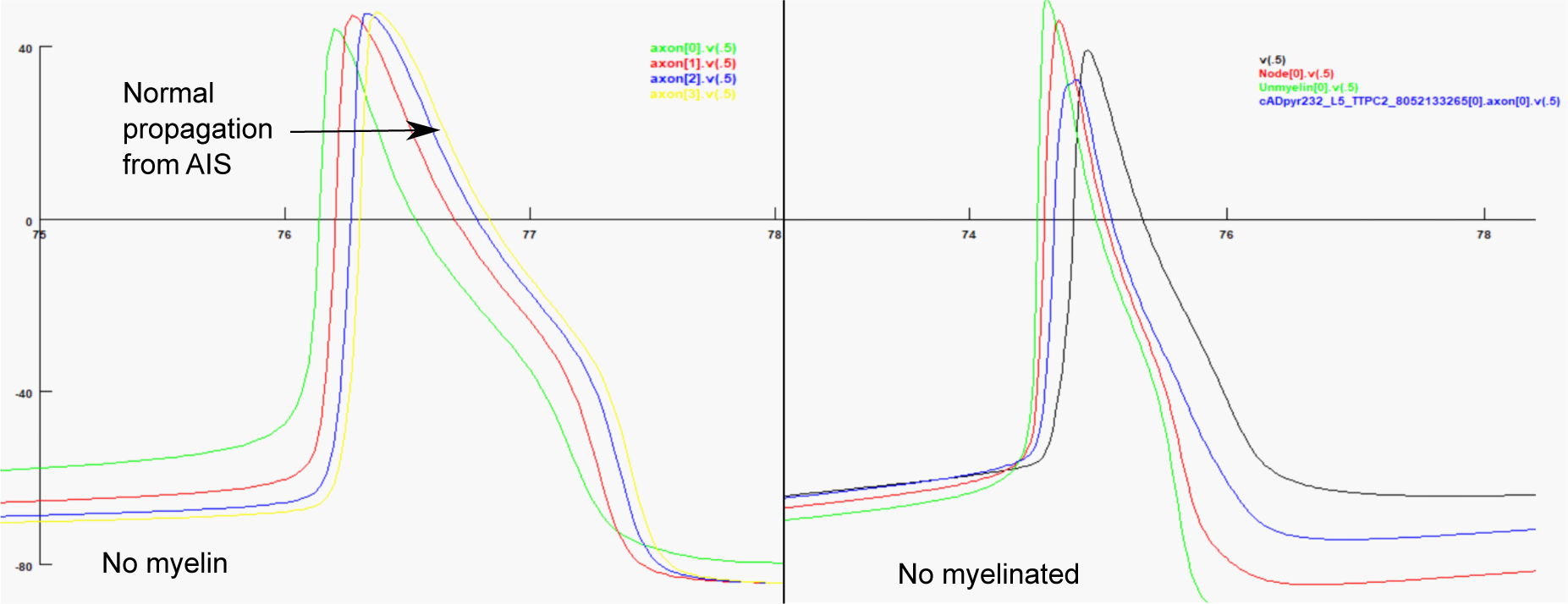
Aberra’s model non-causal problem with an action potential (AP) not propagating from AIS. The left panel illustrates normal behavior when the myelin mechanism is removed, AP propagates from AIS to the axonal tree. In the right panel, the myelinated model shows AP initiation elsewhere on the dendritic tree without the presence of EF. Note that the green curve represents the membrane potential of a compartment far from the soma, therefore not being the AIS.

